# The genome of the camphor tree and the genetic and climatic relevance of the top-geoherbalism in this medicinal plant

**DOI:** 10.1101/2021.10.17.464679

**Authors:** Rihong Jiang, Xinlian Chen, Xuezhu Liao, Dan Peng, Xiaoxu Han, Changsan Zhu, Ping Wang, David E. Hufnagel, Cheng Li, Kaixiang Li, Li Wang

## Abstract

Camphor tree (*Cinnamomum camphora* (L.) J. Presl), a species in the magnoliid family Lauraceae, is known for its rich volatile oils and is used as a medical cardiotonic and as a scent in many perfumed hygiene products. Here, we present a high-quality chromosome-scale genome of *C. camphora* with a scaffold N50 of 64.34 Mb and an assembled genome size of 755.41 Mb. Phylogenetic inference revealed that the magnoliids are a sister group to the clade of eudicots and monocots. Comparative genomic analyses identified two rounds of ancient whole-genome duplication (WGD). Tandem duplicated genes exhibited a higher evolutionary rate, a more recent evolutionary history and a more clustered distribution on chromosomes, contributing to the production of secondary metabolites, especially monoterpenes and sesquiterpenes, which are the principal essential oil components. Three-dimensional analyses of the volatile metabolites, gene expression and climate data of samples with the same genotype grown in different locations showed that low temperature and low precipitation during the cold season modulate the expression of genes in the terpenoid biosynthesis pathways, especially *TPS* genes, which facilitates the accumulation of volatile compounds. Our study lays a theoretical foundation for policy-making regarding the agroforestry applications of camphor tree.

## Introduction

Top-geoherbalism, also known as “Daodi” in China and “Provenance” or “Terroir” in Europe, refers to traditional herbs grown in certain native ranges with better quality and efficacy than those grown elsewhere, in which the relevant characteristics are selected and shaped by thousands of years of the clinical application of traditional medicine (Brinckmann, 2015). The concept of top-geoherbalism is documented in the most ancient and classic Chinese Materia Medica (Divine Husbandman’s Classic of Materia Medica; *Shen Nong Ben Cao Jing*) from approximately 221 B.C. to 220 A.D. (Zhao et al., 2012), which reported that the origin and growing conditions of most herbs were linked to their quality. Historical literature documented cases where the misuse of traditional Chinese medicine (TCM) in prescriptions led to a reduction or absence of therapeutic effects. For example, the application of dried tender shoots of *Cinnamomum cassia* Presl (a component of *Guizhi soup* recorded in *Prescriptions for Emergencies*) from non-top-geoherb regions led to a deficiency in the treatment of fever, while the replacement of this TCM with materials from top-geoherb regions results in proper fever treatment. Currently, TCM from top-geoherb regions accounts for 80% of the market occupancy and economic profits of all TCMs (Huang, 2012), and the significance of top-geoherbs has been revived by the increasing trend of the protection of botanicals with “geographical indication” (GI) (Brinckmann, 2015). Thus, a deep understanding of the pattern and mechanism of top-geoherbalism will strongly guide producers and consumers of TCM and improve the standardization and internalization of the TCM market, especially in the economic context of the Belt and Road Initiative (Hinsley et al., 2020).

A substantial basis of the top-geoherbalism of TCMs is secondary metabolites that play principal roles in the therapeutic effects of herbs. The content of secondary metabolites is a continuous quantitative trait that is determined by three factors: genotype, environment and the interaction between the genotype and environment. Li et al. (2020) showed that cultivars of opium poppy with similar copy number variations in benzylisoquinoline alkaloid biosynthetic genes were likely to exhibit similar alkaloid contents. To identify the ecological and climatic factors driving the development top-geoherbs, the appropriate strategy is to grow herbs with the same genotype (eliminating the effect of genetic variation) in different environmental settings and study how environmental factors are correlated with secondary metabolites by altering gene expression. However, most published studies confound the effects of genetic variation and environmental factors. For example, Tan et al. (2015) identified metabolite markers for distinguishing Radix *Angelica sinensis* from top-geoherb regions and non-top-geoherb regions by analysing the volatile metabolites of processed herbal medicine from drug stores in different regions. The roots of *Paeonia veitchii* showed higher bioactivities when grown at lower average annual temperatures and high elevations based on the analyses of environmental factors and phytochemicals of different samples from seven populations (Yuan et al., 2020). Liu et al. (2020) revealed a correlation between iridoid accumulation and increased temperature by examining 441 individuals from 45 different origins. The confounding factor of genetic variation led to the misinterpretation of how environmental factors alone affect secondary metabolites. In addition, top-geoherb TCMs have been annotated with their cultural properties, including their cultivation, harvesting and postharvest processing (Huang, 2012), which is outside of the scope of the current study.

*Cinnamomum camphora* (L.) J. Presl (Figure 1a), also known as camphor tree is native to China, India, Mongolia and Japan (Chen et al., 2017; Yoshida et al., 1969) and was later introduced to Europe and the southern United States (Bottoni et al., 2021; Hamidpour et al., 2012; Ravindran et al., 2004). Camphor trees are divided into five chemotypes according to their dominant volatile oil components as linalool, camphor, eucalyptol, borneol and isonerolidol types (Luo et al., 2021). Their leaves are especially rich in volatile oil components, including monoterpenes, sesquiterpenes and diterpenes (Hou et al., 2020; Tian et al., 2021). Camphor tree has high medicinal, ornamental, ecological and economic value (Chen, 2020; Liu et al., 2019; Zhang et al., 2020). As a TCM, *C. camphora* can be used to treat rheumatism and arthralgia, sores and swelling, skin itching, poisonous insect bites, etc. Among the many chemical components of the species, camphor is used as a cardiotonic, deodorant and stabilizer in medicine, daily-use chemical production and industry, and linalool is most frequently used as a scent in 60% to 80% of perfumed hygiene products and cleaning agents (Consultation, 2021; Eggersdorfer, 2000; Letizia et al., 2003).

**Figure 1.**
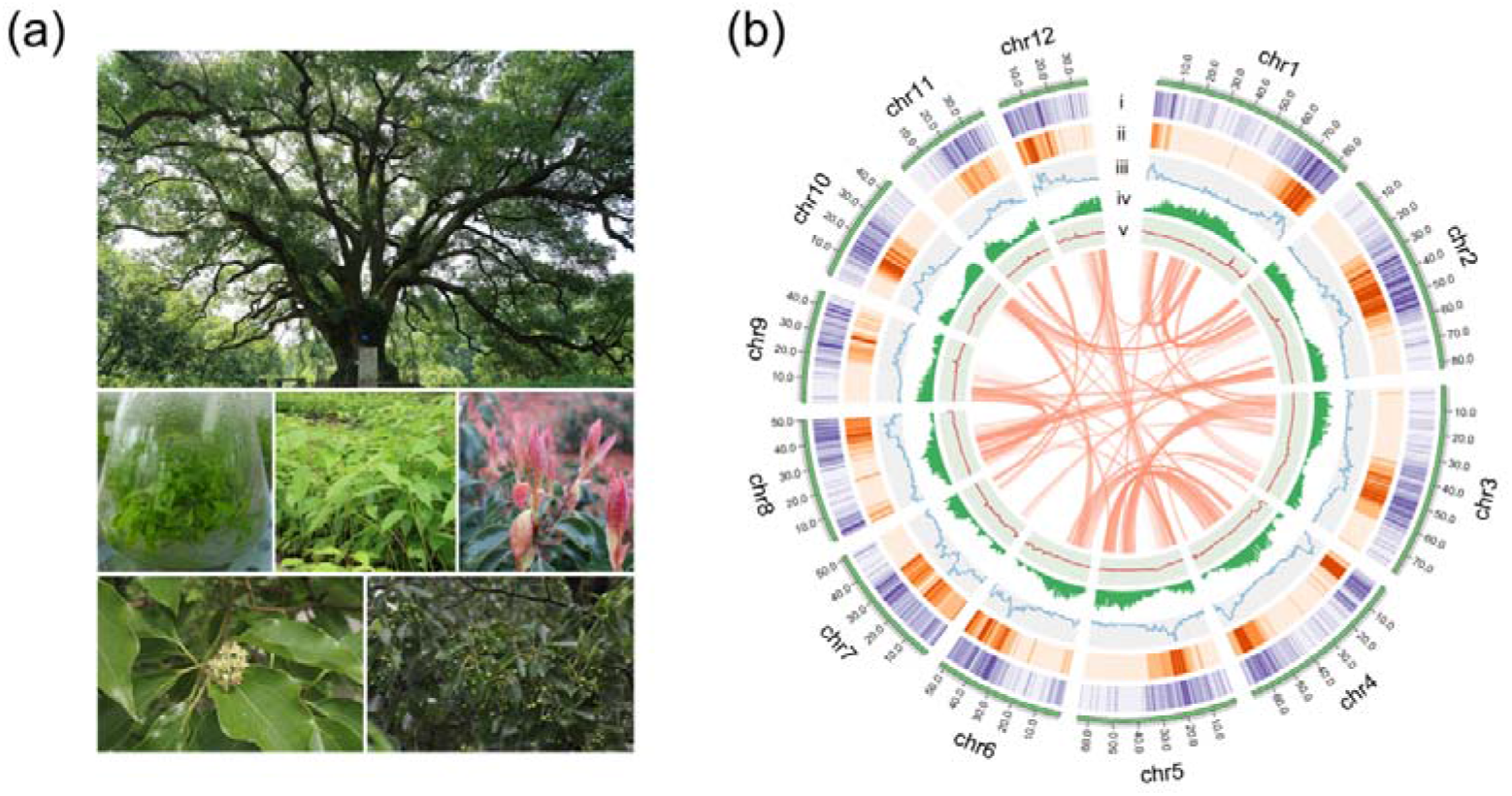
The camphor tree and landscape of its genome. (a) Images of a camphor tree and its multiple tissues. (i) An ancient camphor tree, (ii) tissue-cultured seedings, (iii) seedings, (iv) young leaves, (v) flowers and (vi) fruits. (b) Circos plot of the *C. camphora* genome assembly. Circles from outside to inside: (i) chromosomes, (ii) *Gypsy* LTR density, (iii) *Copia* LTR density, (iii) total LTR density, (iv) gene density and (v) GC content. These density metrics were calculated with 1 Mb nonoverlapped sliding windows. The syntenic genomic blocks (>300 kb) are illustrated with orange lines.

*C. camphora* belongs to Lauraceae within the magnoliid group comprising four orders (Laurales, Magnoliales, Canellales and Piperales). Magnoliids are the third largest group of angiosperms, including approximately 9,000-10,000 species (Massoni et al., 2015; Palmer et al., 2004). From an evolutionary point of view, the mysterious phylogenetic position of magnoliids within angiosperms has been debated for decades. Recently, the debate on the phylogenetic position of magnoliids has focused on three main topologies (Endress and Doyle, 2009; Moore et al., 2007; Qiu et al., 2010; Zeng et al., 2014) positioning magnoliids as (a) a sister group to eudicots (Chaw et al., 2019; Lv et al., 2020; Shang et al., 2020); (b) a sister group to monocots (Qin et al., 2021); or (c) a sister group to the clade of monocots and eudicots (Chen et al., 2019; Chen et al., 2020; Chen et al., 2020; Hu et al., 2019; Rendón-Anaya et al., 2019). Furthermore, the different whole-genome duplication (WGD) events that have occurred in specific lineages of magnoliids and the divergences time between different magnoliid plants remain unclear (Chaw et al., 2019; Chen et al., 2019; Chen et al., 2020; Chen et al., 2020; Hu et al., 2019; Lv et al., 2020; Qin et al., 2021; Rendón-Anaya et al., 2019; Shang et al., 2020).

Despite the economic and evolutionary value of *C. camphora,* the lack of a high-quality genome for the species has greatly restricted the progress of genetic research and the identification of the biosynthetic genes underlying the production of essential volatile compounds with medicinal effects (Chaw et al., 2019; Chen et al., 2019; Chen et al., 2020; Chen et al., 2020; Hu et al., 2019; Lv et al., 2020; Rendón-Anaya et al., 2019; Shang et al., 2020). In our study, we *de novo* assembled the chromosome-scale genome of *C. camphora,* explored the genomic characteristics of *C. camphora* and investigated the genetic and climatic factors underlying the top-geoherbalism of this well-known TCM.

## Results

### *C. camphora* genome assembly and annotation

A haplotype-resolved genome assembly was obtained by using Hifiasm (Cheng et al., 2021) integrating HiFi long reads (24.95 Gb; ~32.83X based on a previous flow cytometry-based estimated genome size of 760 Mb (Wu, 2014)) and Hi-C short reads (63.78 Gb; ~83.92X) with the default parameters. The two contig-level haplotype genomes showed total sizes of 768.97 Mb and 752.05 Mb, respectively, after removing redundant sequences with Purge_haplotigs (Roach et al., 2018). Compared with the earlier published *C. kanehirae* genome (Chaw et al., 2019), the contig N50 was 2.01 Mb for haplotype A and 2.25 Mb for haplotype B; these values are ~2.2- and ~2.5-fold higher, respectively, than the values of the previously published *C. kanehirae* genome (Chaw et al., 2019), respectively (Table 1). Benchmarking Universal Single-Copy Orthologue (BUSCO) analyses showed that the contig sets of haplotype A and haplotype B contained 94.1% and 96.0% complete sets of the core orthologous genes of viridiplantae, respectively (Table S2). Subsequently, the two nonredundant contig sets were anchored to 12 chromosomes based on Hi-C contacts (Figure S1). Overall, the assembled genome size was consistent with previous reports (Table 1; Table S3). As the chromosome-level haplotype A genome was much closer to the estimated genome size than the haplotype B genome, it was used for subsequent analyses.

**Table 1.** The statistics for genome assembles of *Cinnamomum*

|  | *C. camphora* | *C. kanehirae* |
| --- | --- | --- |
| **Gene assembly** |  |  |
| Genome size (Mb) | 755.41 | 730.43 |
| Scaffold number (n) | 1,037 | 2,153 |
| Scaffold N50 (Mb) | 64.34 | 50.35 |
| Scaffold L51 (n) | 5 | 6 |
| Chromosome-scale scaffolds (Mb) | 697.81 (92.38%) | 672.85 (92.12%) |
| Contig number (n) | 1,686 | - |
| Contig N50 (Mb) | 2.01 | 0.90 |
| Contig L51 (n) | 108 | - |
| GC content of the genome (%) | 39.49 | 38.22 |
| Complete BUSCOs (%) | 96.20 | 88.50 |
| **TE annotation** |  |  |
| TE content (%) | 47.86 | 47.87 |
| LTR content (%) | 27.94 | 25.53 |
| Copia content (%) | 7.69 | - |
| Gypsy content (%) | 15.42 | - |
| LAI | 13.82 | - |
| **Gene annotation** |  |  |
| Number of predicted genes (n) | 24,883 | 27,899 |
| Average gene length (bp) | 9,295.00 | 7,591.84 |
| Average CDS length (bp) | 1,189.59 | 1,310.51 |
| Average exon length (bp) | 309.29 | 241.55 |
| Average exon number per transcript (n) | 4.80 | 5.40 |
| Average intron length (Mb) | 1,548.48 | 1,425.80 |
| Average intron number per transcript (n) | 4.40 | 4.06 |
| **Gene function annotation** |  |  |
| Nr | 24,147 (97.04%) | - |
| Swissprot | 20,122 (80.87%) | - |
| Pfam | 17,945 (72.12%) | - |
| GO | 11,593 (46.59%) | - |
| KEGG | 11,218 (45.08%) | - |
| Total | 24,152 (97.06%) | - |

The assembly for *C. camphora* was compared with *C. kanehirae*. Dash (-) indicates data were not shown in the original research.

The final assembled *C. camphora* genome consisted of 12 chromosomes, 1,025 scaffolds and 1,686 contigs, with a scaffold N50 of 64.34 Mb (Table 1; Figure 1b). The number of assembled chromosomes was consistent with previous cytological observations (http://ccdb.tau.ac.il). The length of the chromosomes ranged from 84.54 Mb (Chr1) to 36.36 Mb (Chr12) (Table S5). The high fidelity of the assembly was corroborated by multiple lines of evidence. First, the mapping rates of RNA-Seq paired-end reads against the assembled genome were high (93%-95%). Second, the high completeness of this assembly was supported by a 96.2% BUSCO value (Table 1; Table S4), suggesting high completeness at the gene level, which was much better than the completeness of the *C. kanehirae* genome assembled via PacBio CLR sequencing. Third, the long terminal repeat (LTR) assembly index (LAI) score (Ou et al., 2018) was 13.82, indicating the “reference” level of the genome and reflecting completeness at the transposable element (TE) level.

By combining *ab initio* prediction, orthologous protein and transcriptomic data, we annotated 24,883 protein-coding genes using the MAKER pipeline (Cantarel et al., 2008) (Table 1). The average lengths of the gene regions, coding sequences (CDSs), exon sequences, and intron sequences were 9,295.00, 1,189.59, 309.29, and 1,548.48 bp, respectively (Table 1). Among the predicted protein-coding genes, 97.06% could be annotated in at least one of the following protein-related databases: NCBI nonredundant protein (NR) (97.04%), Swiss-Prot (80.87%), Pfam (72.12%), GO (46.59%) or KEGG (45.08%) (Table 1).

A total of 361.37 Mb of TEs were identified (Table S6). Long-terminal repeat retrotransposons (LTR-RTs) were the most abundant type of repetitive sequence, accounting for 46.86% of the whole genome, similar to the proportion of LTRs found in *C. kanehirae* (Table 1). A total of 3,731 intact LTR-RTs were identified, and the frequency distribution of insertion times showed a burst of LTR-RTs 1-2 million years ago (Mya) (Figure S2), consistent with previous reports in magnoliids. Gypsy elements were the largest LTR-RT superfamily in *C. camphora,* constituting 15.42% of the genome. The second largest superfamily was Copia, accounting for 7.69% of the genome. Other unclassified LTR-RTs encompassed 4.83% of the genome. Among DNA retrotransposons, terminal inverted repeat sequences (TIRs) and non-TIRs comprised 14.05% and 5.87% of the genome, respectively.

### Phylogenetic and WGD analyses

The concatenate phylogenetic tree constructed with 172 single-copy orthologous genes among 16 species showed that *C. camphora* was clustered with two other Lauraceae species and that the formed clade was then clustered with other magnoliids (Figure 2; Figure S3a). More importantly, the magnoliids were sister to the combined clade of eudicots and monocots rather than to either monocots or eudicots. In addition, the coalescence-based phylogenetic tree showed the same topology for magnoliids as the concatenation tree (Figure S3b). This phylogenetic topology is consistent with some previous studies (Chen et al., 2019; Chen et al., 2020; Chen et al., 2020; Hu et al., 2019; Rendón-Anaya et al., 2019) but contrasts with other reports (Chaw et al., 2019; Lv et al., 2020; Qin et al., 2021; Shang et al., 2020). The divergence time between magnoliids and the clade of monocots and eudicots was inferred to be 147.47-170.11 Mya (Figure 2) in MCMCTree with fossil calibration (Yang et al., 2007), coinciding with the estimates in other studies (Chen et al., 2019; Chen et al., 2020; Chen et al., 2020; Hu et al., 2019; Rendón-Anaya et al., 2019). In the clade of magnoliids, the divergence time between Laurales and Magnoliales was approximately 138.96 Mya, after which the divergence of Lauraceae and Calycanthaceae occurred approximately 114.26 Mya, and finally, *C. camphora* diverged from the most recent common ancestor of *C. camphora* and *C. kanehirae* approximately 28.03 Mya.

**Figure 2.**
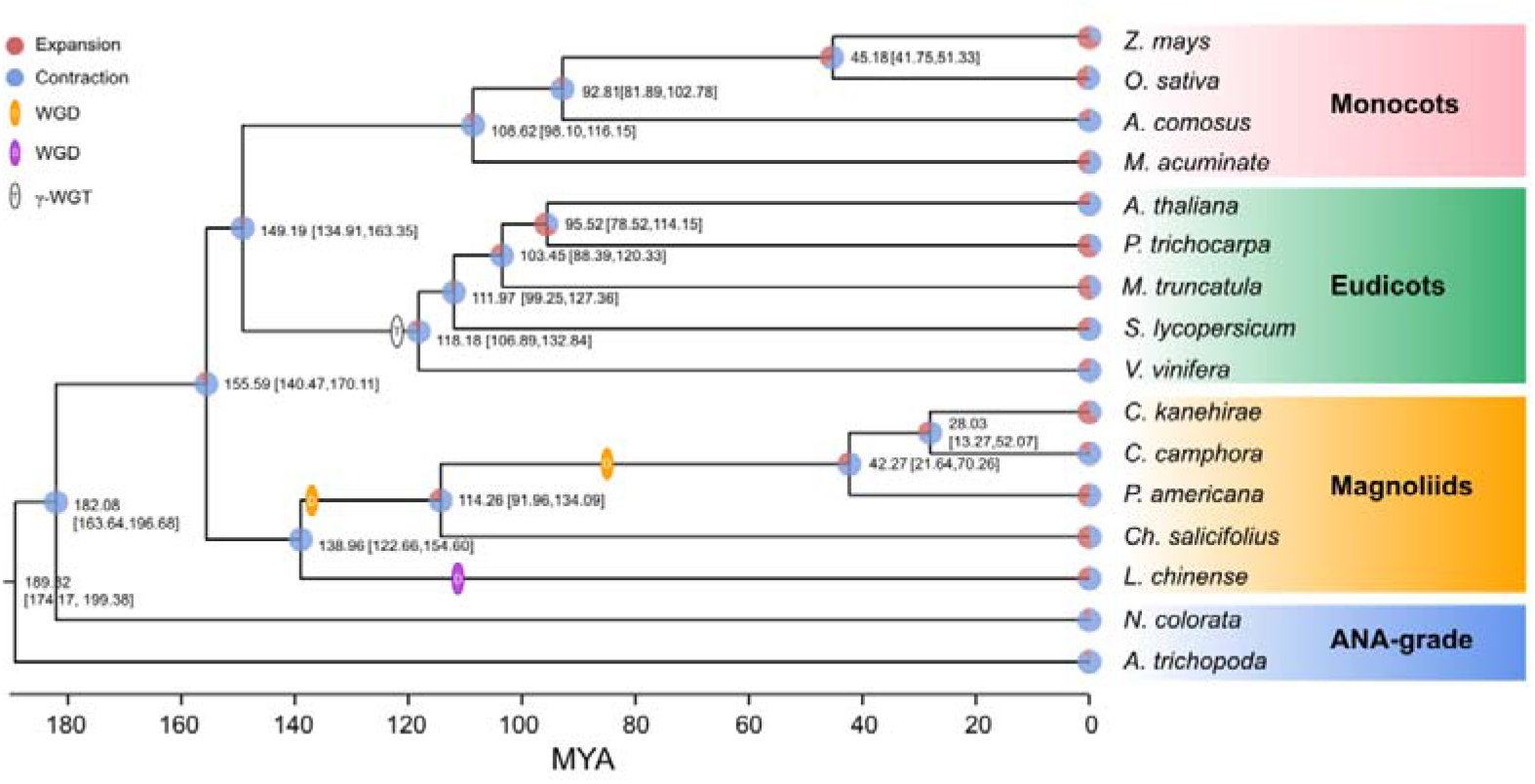
Phylogenetic analyses. The phylogenetic tree was constructed based on 172 single-copy orthologous genes of 16 species using two ANA-grade species as outgroups; node age and 95% confidence intervals are labeled. Pie charts show the proportions of gene families that underwent expansion or contraction. Predicted whole-genome duplication (WGD) events were only indicated for Laurales and Magnoliales.

The intragenomic collinearity and syntenic depth ratio of *C. camphora* showed clear syntenic evidence of ancient WGD events (Figure 1b; Figure S4). Only the paralogous gene set derived from duplicates produced by WGD was used for the analyses of synonymous substitutions per site (*Ks*) (see “Methods” section). Two ancient WGD events shared within the Laurales lineage (*C. camphora*, *C. kanehirae* and *Persea americana*) occurred approximately 85.66 Mya and 137.90 Mya, represented by two signature peaks with *Ks* values of approximately 0.52 and 0.83 (Figure 3a). The recent *Ks* peak (85.66 Mya) occurred much earlier than the divergence time (42.27 Mya) between *P. americana* and the common ancestor of *Cinnamomum*, indicating that this round of WGD was shared between *Cinnamomum* and *Persea* (Figure 2; Figure 3a). Additionally, the *Ks* peak (137.90 Mya) occurred earlier than the divergence (114.26 Mya) between Lauraceae species and *Chimonanthus salicifolius*, implying that this round of WGD was shared between Lauraceae and Calycanthaceae (Figure 2; Figure 3a). We also detected a WGD event in *Liriodendron chinense* that occurred approximately 116.67 Mya, corresponding to a *Ks* peak of 0.70 (Figure 3a), which is consistent with a previous study (Chen et al., 2019).

**Figure 3.**
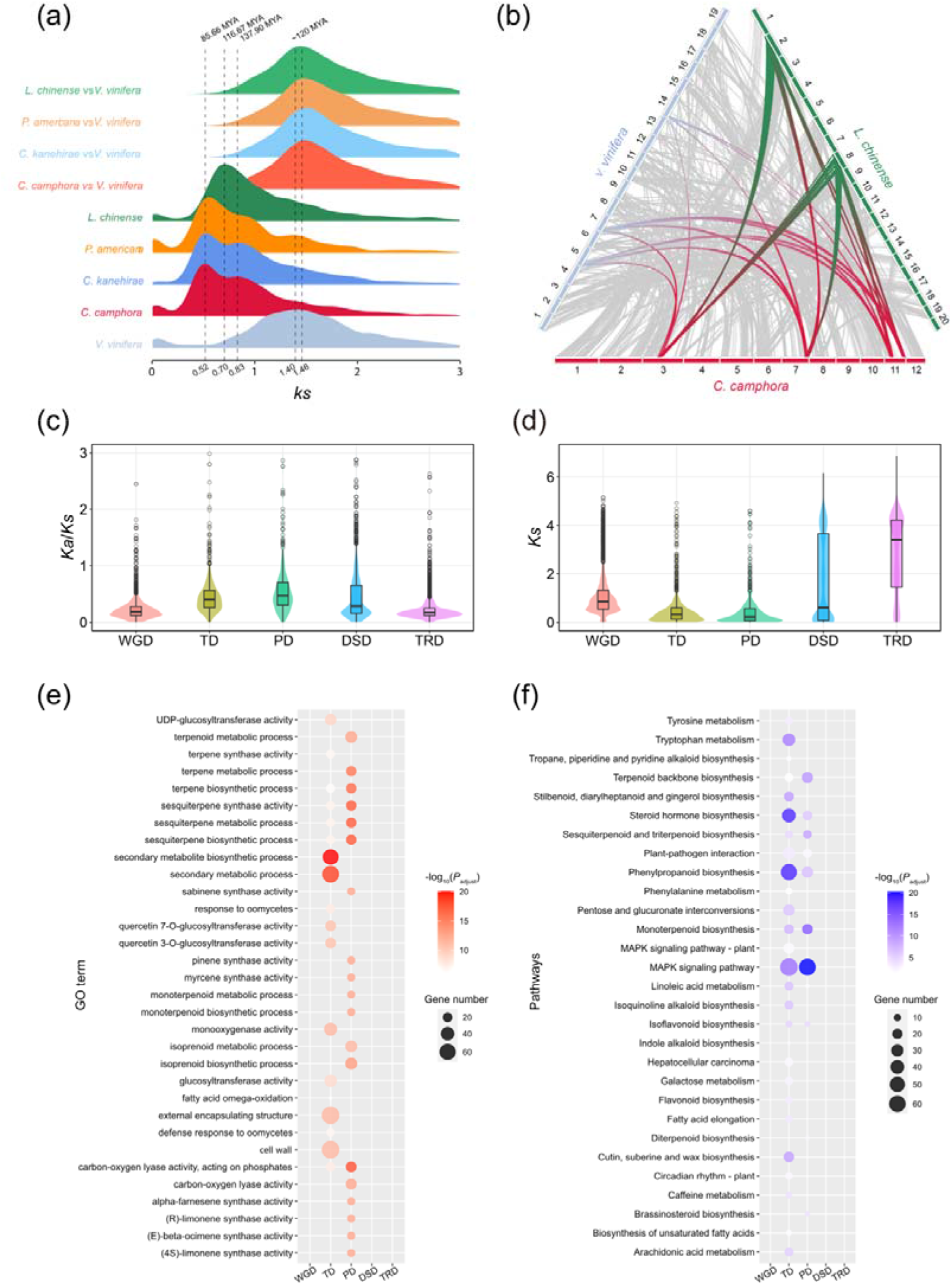
Gene duplication and evolution. (a) *Ks* distribution of paralogues in magnoliid species (*C. camphora*, *C. kanehirae*, *P. americana* and *L. chinense*) and orthologues between these magnoliids and *V. vinifera*. (b) Synteny blocks among *C. camphora*, *L. chinense* and *V. vinifera*. (c) *Ka/Ks* ratio distributions of gene pairs derived from different types of duplication. WGD, whole-genome duplication; TD, tandem duplication; PD, proximal duplication; TRD, transposed duplication; DSD, dispersed duplication. (d) *Ks* ratio distributions of gene pairs derived from different types of duplication. (e) GO enrichment analyses of genes from different types of duplication. The enriched GO terms with adjusted *P* values < 0.01 are presented. The colours of the bubbles indicate the statistical significance of the enriched GO terms. The sizes of the bubbles indicate the number of genes associated with one GO term. (f) KEGG enrichment analyses of genes resulting from different types of duplication. Enriched KEGG pathways with adjusted *P* values < 0.01 are presented. The colours of the bubbles represent the statistical significance of enriched KEGG pathways. The sizes of the bubbles indicate the number of genes associated with one KEGG pathway.

To confirm that *C. camphora* underwent two rounds of ancient WGD, we performed intergenomic synteny analyses between the genomes of *C. camphora* and *L. chinense* together with *Vitis vinifera*. A “4:2” syntenic relationship was detected between *C. camphora* and *L. chinense* (Figure S5a). Given the hexaploidy event (γ-WGT) shared among core eudicots, including *V. vinifera* (French-Italian Public Consortium for Grapevine Genome Characterization, 2007), an overall “4:3” syntenic relationship was observed between *C. camphora* and *V. vinifera* (Figure S5b). Furthermore, we discovered a “2:3” syntenic relationship between *L. chinense* and *V. vinifera* (Figure S5c). Thus, the syntenic relationships among the *L. chinense:V*. *vinifera:C. camphora* genomes showed a 2:3:4 ratio (Figure 3b). Our results echoed previous studies (Chaw et al., 2019; Chen et al., 2019), indicating two rounds of ancient WGD in Lauraceae.

To determine the functional roles of the genes retained after WGD events, GO and KEGG enrichment analyses were performed. The results showed that the genes retained after WGD were enriched in basic physiological activities, processes and pathways, such as structural constituents of ribosomes, NADPH dehydrogenase activity, photosynthesis and plant hormone signal transduction, suggesting that two rounds of WGD events enhanced the adaptability of *C. camphora* to changing environments by improving basic physiological activities and primary metabolism (Figure S6; Figure S7; Table S7; Table S12).

### Tandem and proximal duplications contribute to terpene synthesis in *C. camphora*

A total of 18,938 duplicated genes were identified, including 7,043 WGD genes (37.19%), 1,597 tandem duplication (TD) genes (8.43%), 1,079 proximal duplication (PD) genes (5.70%), 5,127 dispersed duplication (DSD) genes (27.07%) and 3,779 transposed duplication (TRD) genes (19.95%) (Figure S8). The *Ks* and *Ka*/*Ks* values among different types of duplicated genes were calculated (Figure 3c; Figure 3d). The *Ks* values of TD and PD genes were much smaller than those of other types of duplications (Figure 3d), suggesting that TD and PD genes were formed recently. Additionally, the higher *Ka*/*Ks* ratios of TD and PD genes (Figure 3c) implies that they were subject to more rapid sequence divergence.

We also performed GO and KEGG enrichment analyses of the gene sets associated with the five duplication types (Figure S6; Figure S7; Table S7-S16). The GO results showed that the biological process (BP) and molecular function (MF) categories of secondary metabolite, monoterpene, sesquiterpene and terpene biosynthesis and metabolism were significantly enriched in the TD and PD gene sets, but no enrichment of these GO terms was found in the WGD, DSD and TRD gene sets (Figure 3e). Regarding the KEGG results, the TD and PD gene sets were significantly enriched in the terpenoid backbone, monoterpenoid, sesquiterpenoid and phenylpropanoid biosynthesis pathways. These secondary metabolite biosynthesis pathways were not enriched in the other three duplication gene sets (Figure 3f). In addition, KEGG pathways related to responses to environmental stimuli, such as the MAPK signalling pathway, steroid hormone biosynthesis and plant-pathogen interaction, were also significantly enriched in the TD and PD gene sets. All of these results indicated that the rapidly evolving TD and PD genes played essential roles in the synthesis of secondary metabolites, especially monoterpenes and sesquiterpenes, and the response to environmental stimuli in *C. camphora*.

### Metabolic reflection of top-geoherbalism

To evaluate how environmental factors affect metabolite accumulation, we grew tissue-cultured seedlings from a single mother plant in four major production locations, including Qinzhou, Nanning, Baise and Liuzhou, beginning in May 2018. Mature leaves were harvested from the four locations in November 2020 for RNA-Seq and metabolite assessment. Thus, we were able to compare the effects of environmental conditions on metabolite accumulation based on the fixed genotype.

Volatile metabolites are a substantial basis of the top-geoherbalism of camphor tree. Based on targeted gas chromatography-mass spectrometry (GC-MS) analyses, we identified 153 nonredundant volatile metabolites (Supplemental Table 17). The 153 volatile metabolites included 82 terpenes (53.60%), 14 esters (9.15%), 13 ketones (8.50%), 12 heterocyclic compounds (7.84%), 10 alcohols (6.54%), 9 aromatics (5.88%) and 13 additional metabolites (8.49%) that did not belong to these six main types (Figure 4a). For quality control, we performed hierarchical clustering and principal component analyses (PCA) of metabolic abundances in these 24 samples (Figure S9). The first replicate in Baise (“Baise_1”) was identified as an outlier and was removed from further analyses.

**Figure 4.**
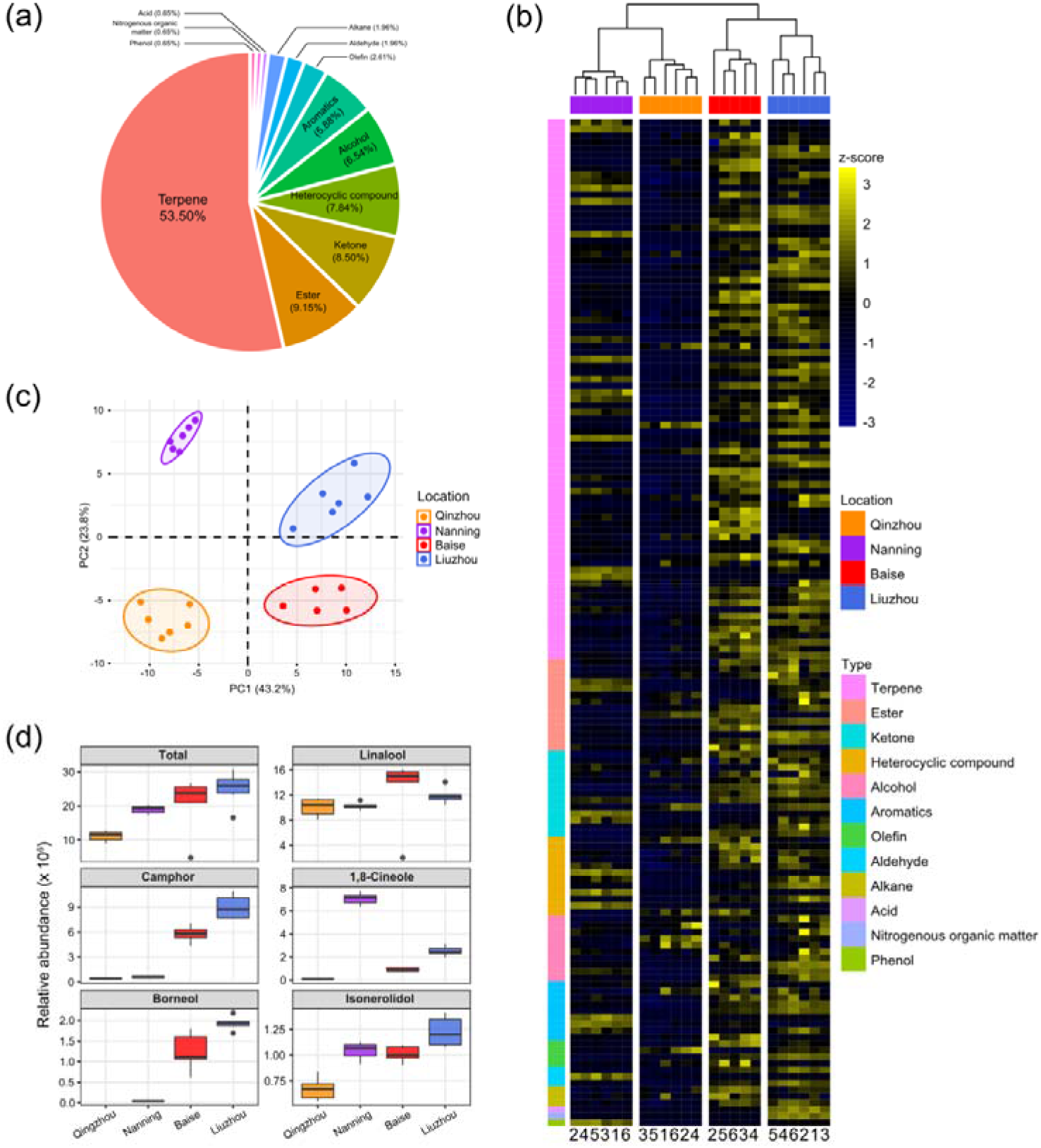
Abundance patterns of volatile metabolites in *C. camphora* planted in different locations. (a) Pie charts show the proportions of different types of metabolites identified in the current study. (b) Hierarchical clustering heatmap of metabolic abundance profiles in the four planting locations, including Qinzhou, Nanning, Baise and Liuzhou, indicated on the x axis. Metabolic abundance was averaged and z-score transformed. The rows are clustered by the types of metabolites. (c) Principal component analyses (PCA) of metabolites of *C. camphora* planted in the four locations. The circles represent the 95% confidence intervals. (d) The relative abundances of linalool, borneol, camphor, 1,8-cineole and isonerolidol in the four planting locations.

The volatile metabolites of the samples from the four locations exhibited distinct clustering in the PCA plot (Figure 4c). Notably, the metabolites of samples from Baise and Liuzhou were clearly separated from those of Qinzhou and Nanning according to principal component 1 (PC1), which accounted for 43.2% of the total variance. The heatmap based on the relative abundances of volatile metabolites showed that samples from Baise and Liuzhou exhibited higher abundances than those from Qinzhou and Nanning, especially with regard to terpenes (Figure 4b; Figure S10).

Linalool, borneol, camphor, 1,8-cineole and isonerolidol are the five main phytochemicals determining the medicinal value and top-geoherbalism of *C. camphora*. A detailed comparison of the five volatile terpenoids implied that samples from Liuzhou showed the highest total abundance, followed by the samples from Baise, while the abundance was lowest in Qinzhou (Figure 4d). The abundance profiling of camphor and borneol in the four locations echoed the trends of the total abundance of the five volatile terpenoids. The abundance of linalool and isonerolidol peaked in Baise and Liuzhou, respectively, while that of 1,8-cineole peaked in Nanning. Taken together, these results suggested that Baise and Liuzhou are top-geoherb regions of the camphor tree in Guangxi Province.

### Modulation of gene expression related to top-geoherbalism

TPSs are the rate-limiting enzymes in the production of terpenoids (Chen et al., 2011; Tholl, 2015), including monoterpenes, sesquiterpenes and diterpenoids (see “Methods” section). The identified *TPS* genes of *C. camphora*, *C. kanehirae*, *P. americana*, *L. chinense* and *Arabidopsis thaliana* were clustered into six clades in the phylogenetic tree, corresponding to the *TPS-a*, *TPS-b*, *TPS-c*, *TPS-e*, *TPS-f* and *TPS-g* subfamilies (Chen et al., 2011), among which *TPS-b* and *TPS-g* encode the enzymes catalysing the production of 10-carbon monoterpenoids from geranyl diphosphate (GPP), and *TPS-a* genes are responsible for catalysing the production of 15-carbon sesquiterpenoids from farnesyl diphosphate (FPP) (Figure 5a,b). The copy number variation of the *TPS* genes showed that *TPS-b* was greatly expanded in *C. kanehirae* and *C. camphora* (Figure 5a,b), which resulted in the high abundance of monoterpenoids in *Cinnamomum* (Chen et al., 2020; Chen et al., 2020). Specifically, 44 *TPS-b* genes were identified in *C. camphora*, accounting for 61% of all its *TPS* genes, consistent with the percentage observed in *C. kanehirae* (63%) and much higher than those in *L. chinense* (33%), *A. thaliana* (18%) and *P. americana* (12%). We also observed more copies of the *TPS-g* subfamily in *C. camphora* than in *A. thaliana* and *P. americana*. However, no expansion of *TPS-a* genes was detected in *Cinnamomum*.

**Figure 5.**
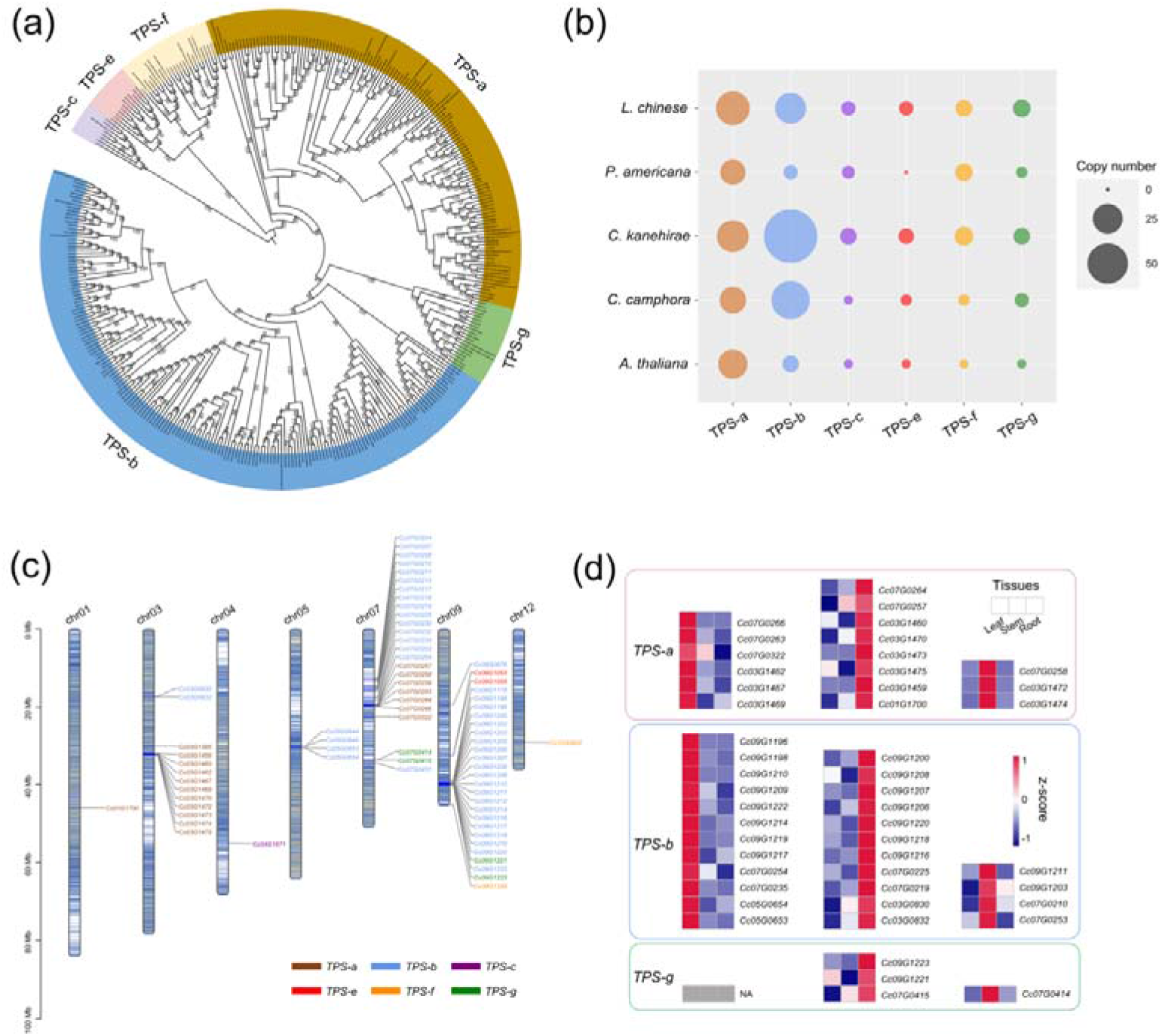
Genes involved in the biosynthesis of volatile terpenoids. (a) Phylogenetic analyses of *TPS* genes in *C. camphora*. The phylogenetic tree was constructed based on *TPS* gene sequences from four magnoliid genomes (*C. camphora*, *C. kanehirae*, *P. americana* and *L. chinense*) and *A. thaliana*. (b) Copy numbers of *TPS* genes in the genomes of four magnoliids and *A. thaliana*. (c) Distribution of the *TPS* genes on seven chromosomes of *C. camphora*. (d) Tissue-specific expressions of *TPS-a*, *TPS-b* and *TPS-g* subfamilies.

Next, we examined how the *TPS* genes were distributed across the genome. Chromosome 9 and chromosome 7 harboured the most *TPS* genes (27 and 25, respectively), followed by chromosome 3 (13 *TPS* genes) (Figure 5c; Table S18). Interestingly, genes from the seven subfamilies were observed as tandem duplicates. Two large *TPS-b* gene clusters were identified on chromosome 9 (21 genes; ca. 38.78-40.12 Mb) and chromosome 7 (10 genes; ca. 12.70-16.03 Mb), respectively. In addition, two large *TPS-a* gene clusters were detected on chromosome 3 (10 genes; ca. 31.90-32.52 Mb) and chromosome 7 (6 genes; ca. 19.49-19.90 Mb), respectively. The remaining smaller *TPS* gene clusters were scattered throughout the genome of *C. camphora*.

Notably, the *TPS* genes exhibited a strong tissue-specific expression pattern (Figure 5d; Table S19), especially *TPS-a*, *TPS-b* and *TPS-g*. We downloaded previously published RNA-Seq data (https://www.ncbi.nlm.nih.gov/bioproject/PRJNA747104) from three tissues (leaf, stem and root) of *C. camphora* to determine the gene expression profile of *TPS* genes. As leaves are the major tissue used in medicine, we focused on the *TPS* genes with high expression levels in leaves. Six *TPS-a* genes and twelve *TPS-b* genes showed higher expression in leaves than in stems and roots. All of these results indicated that the expansion of the *TPS-b* subfamily and the tandem duplication of *TPS-b* genes probably contribute to monoterpenoid biosynthesis in *C. camphora*.

The biosynthetic pathways of terpenoids are derived from isopentenyl diphosphate (IPP) and dimethylallyl diphosphate (DMAPP) produced via the methylerythritol phosphate (MEP) and mevalonate (MVA) pathways, respectively (Li et al., 2019; Zhou and Pichersky, 2020). Comparative transcriptome analyses of samples from the four locations were performed to examine the expression patterns of genes involved in the MEP and MVA pathways. We combined four differentially expressed gene (DEG) sets identified in Liuzhou vs. Qinzhou, Liuzhou vs. Nanning, Baise vs. Qinzhou and Baise vs. Nanning (Figure S11). We detected 22 and 23 DEGs in the MEP and MVA pathways, respectively (Figure 6; Table S20). Generally, the DEGs involved in all the steps of the MEP and MVA pathways showed higher expression in Baise and Liuzhou than in Qinzhou and Nanning, except for the *2*-*C*-*methyl*-*D*-*erythritol*-*2*,*4*-*cyclodiphosphate synthase* (*MDS*) gene, which was absent in the combined DEG set. The same expression pattern was observed for the genes related to downstream steps, including *isopentenyl diphosphate isomerase* (*IDI*), *geranyl diphosphate synthetase* (*GPPS*), *farnesyl diphosphate synthetase* (*FPPS*), *TPS-a*, *TPS-b* and *TPS-g*. The modulation of gene expression in the terpenoid biosynthesis pathway echoed the higher accumulation of monoterpenoids and sesquiterpenoids observed in Baise and Liuzhou than in Qinzhou and Nanning.

**Figure 6.**
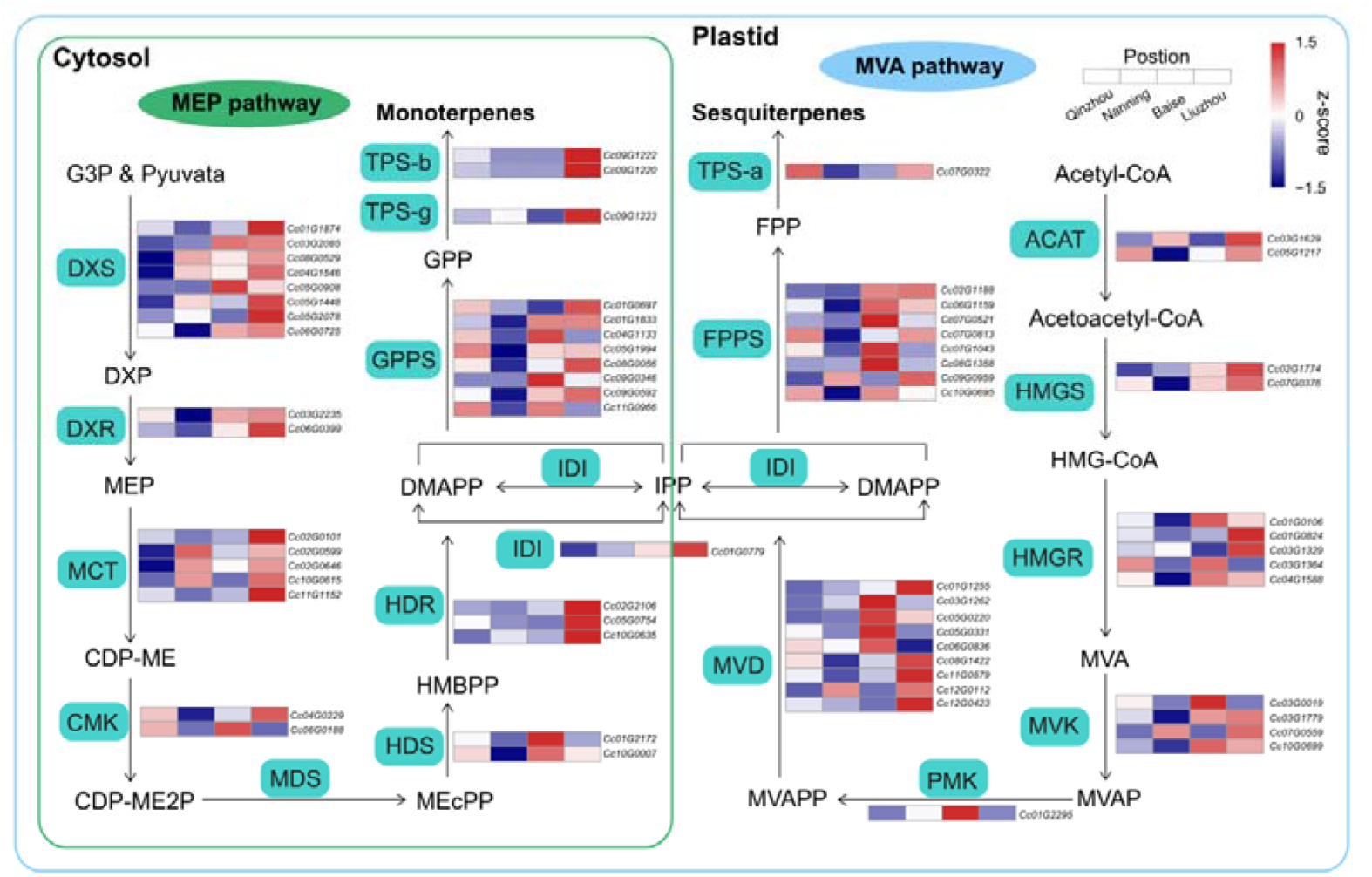
Biosynthetic pathways of monoterpenoids and sesquiterpenoids. Relative expression profiling of genes involved in volatile terpenoid biosynthesis among the four planting locations (Qinzhou, Nanning, Baise and Liuzhou). Gene expression was extracted from the combined differentially expressed gene (DEG) set (Liuzhou vs. Qinzhou, Liuzhou vs. Nanning, Baise vs. Qinzhou and Baise vs. Nanning). MEP, mevalonate pathway; MEP, methylerythritol phosphate pathway; ACAT, acyl-coenzyme A-cholesterol acyl-transferase; HMGS, hydroxymethylglutaryl coenzyme A synthase; HMGR, hydroxymethylglutaryl coenzyme A reductase; MVK, mevalonate kinase; PMK, phospho-mevalonate kinase; MVD, mevalonate diphosphate decarboxylase; DXS, 1-deoxy-D-xylulose 5-phosphate synthase; DXR, 1-deoxy-D-xylulose 5-phosphate reductoisomerase; MCT, 2-C-methyl-D-erythritol-4-phosphate cytidylyltransferase; CMK, 4-(cytidine-5-diphospho)-2-C-methyl-D-erythritol kinase; MDS, 2-C-methyl-D-erythritol-2,4-cyclodiphosphate synthase; HDS, (E)-4-hydroxy-3-methyl-but-2-enyl-pyrophosphate synthase; HDR, (E)-4-hydroxy-3-methyl-but-2-enyl-pyrophosphate reductase.

### Climatic factors underlying the top-geoherbalism of *C. camphora*

To determine what climatic factors caused the differences in volatile metabolites among the *C. camphora* plants of the same genotype grown in different locations (Figure 7a), we downloaded the climate data of Qinzhou, Nanning, Baise and Liuzhou from three years (2018-2020) from the National Meteorological Information Centre (http://data.cma.cn/en). The 17 climatic factors could be classified into temperature-related, wind-related, pressure-related, precipitation-related, humidity-related and sunshine-related factors. (Table S21; Table S22).

**Figure 7.**
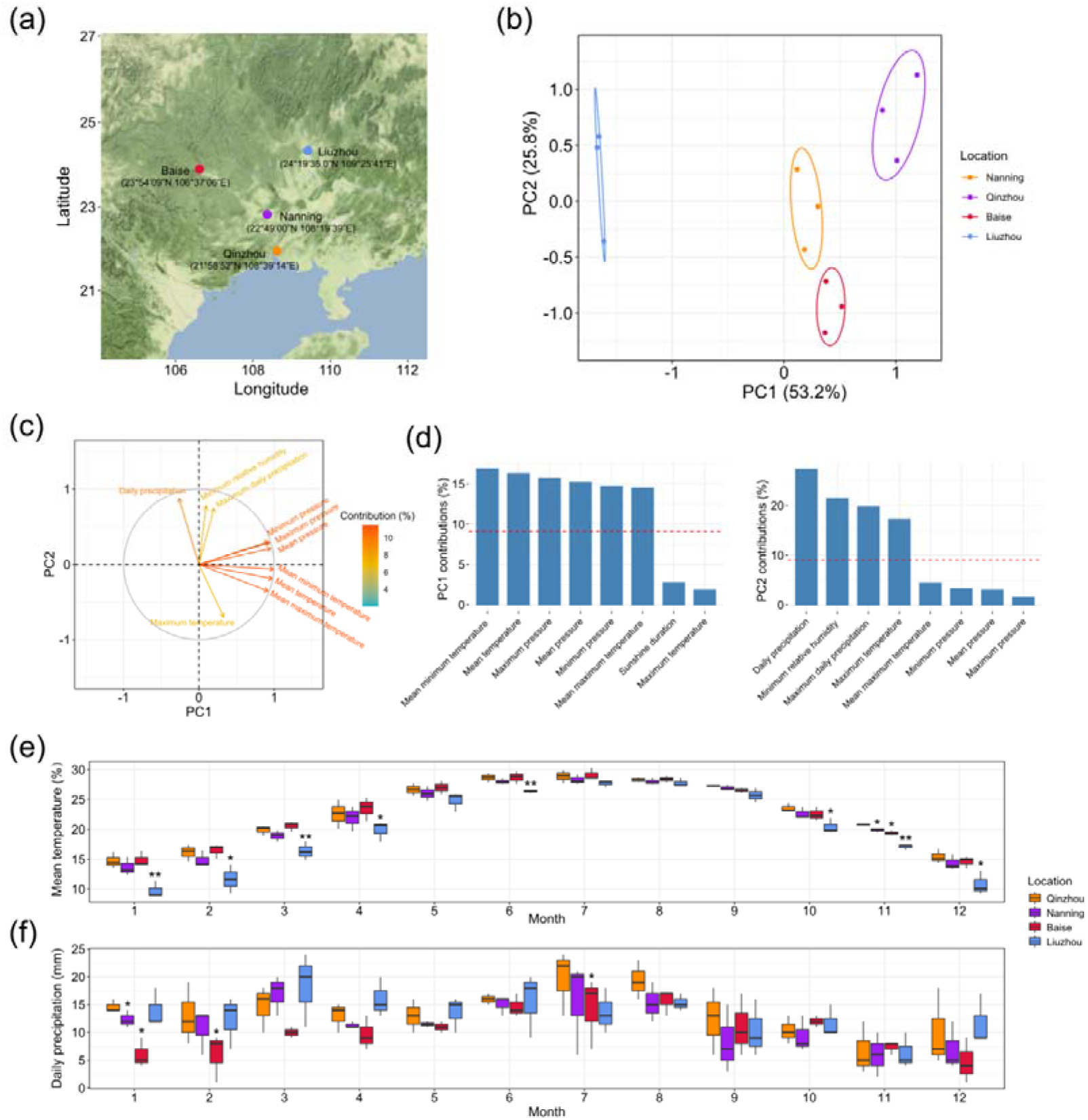
Analyses of climatic factors in different planting locations. (a) Geographical distribution of *C. camphora* planting locations, including Qinzhou, Nanning, Baise and Liuzhou. (b) Principal component analyses (PCA) of seventeen climatic factors in the four planting locations in 2018, 2019 and 2020. (c) The loadings of climatic factors in the PCA plot. The colours of the arrows represent the percentages of the contributions of climatic factors to the PCs. (d) Histograms of the percentages of the contributions of different climatic factors to PC1 and PC2. The red dashed lines indicate the average contributions of different climatic factors. Only the top eight climatic factors are shown. (e) Monthly observations of the mean temperature in the four planting locations. Single and double asterisks indicate statistically significance levels of *P* < 0.05 and *P* < 0.01, respectively, between Liuzhou and Qinzhou/Nanning (paired-sample Student’s *t* test). (f) Monthly observations of daily precipitation in the four planting locations. A single asterisk indicates the statistically significance levels of *P* < 0.05 between Baise and Qinzhou/Nanning (paired-sample Student’s *t* test).

PCAs of the climatic factors showed that PC1 (accounting for 53.2% of the total variance) separated Liuzhou from Qinzhou, Nanning and Baise, while PC2 (accounting for 25.8% of the total variance) split Baise from Qinzhou, Nanning and Liuzhou (Figure 7b). To examine the contributions of climatic factors to PC1 and PC2, we loaded them in the PCA plot (Figure 7c). Temperature-related factors were the main variables contributing to PC1, including mean minimum temperature (16.88%), mean temperature (16.32%) and mean maximum temperature (14.50%), and precipitation-related factors mainly contributed to PC2, including daily precipitation (27.29%) and maximum daily precipitation (19.75%) (Figure 7d).

To examine the detailed differences in temperature- and precipitation-related factors in these four locations, we plotted the monthly observations of mean minimum temperature, mean temperature, mean maximum temperature, daily precipitation and maximum daily precipitation (main contributors to PC1 and PC2) from 2018 to 2020 (Figure 7e, f; Figure S12; Table S22). The mean temperature of Liuzhou was much lower than those of the other locations, especially in the two colder quarters of the year. Additionally, daily precipitation was lower in Baise than in the other locations, especially in early spring. All of these results suggested that relatively low temperature and precipitation during the cold season imposes some degree of stress on the plants and thus stimulated the production of the desired terpenoids in *C. camphora*.

## Discussion

Here, by decoding the genome of the camphor tree, we revealed that magnoliids are a sister group to the clade of eudicots and monocots. The two rounds of WGD identified in *C. camphora* were dated to ca. 85.66 Mya and 137.90 Mya, respectively. The former was shared with Lauraceae species, and the latter occurred before the divergence of Lauraceae and Calycanthaceae. We found that rapidly evolving TD genes play key roles in the synthesis of secondary metabolites, especially for monoterpenes and sesquiterpenes. By analysing volatile metabolites in leaves sampled from plants of the same genotype grown in four different locations, we found higher accumulation of the key volatile metabolites in regions with lower temperature and precipitation in the cold season, which was attributed to the differential expression of genes related to the MVA and MEP pathways as well as their downstream steps in terpene biosynthesis. Our study confirmed that abiotic stress contributes to the development of top-geoherbs, laying a theoretical foundation for policy making on the agroforestry applications of camphor trees.

In addition to releasing the first chromosome-level genome of this significant economic tree species, our study is innovative in two regards. First, it is among the very few innovative three-dimensional evaluations of the top-geoherbalism of a TCM species, integrating phytochemical, genetic and climatic analyses of camphor tree germplasm (Li et al., 2017). Second, our study is the first to fix the genotypic variation of the germplasm so that it is feasible to investigate how climatic factors alone affect the accumulation of metabolites (Li et al., 2020; Mandim et al., 2021). By integrating these two novel approaches, our study provides an example of how to comprehensively scrutinize the genetic and climatic factors affecting the composition of secondary metabolites.

The dominant climatic factors affecting the secondary metabolites of the camphor tree are not always the key factors affecting in other traditional herbs. For example, humidity and sunshine time are the chief limiting factors in the production of artemisinin in *Artemisia annua* L. (Li et al., 2017). Additionally, lower temperature in the coldest quarter imposes stress on camphor tree and increases the production of desired volatile compounds. In other cases, a higher temperature imposes stress and enhances the production of desired metabolites (Liu et al., 2020). The limiting ecological and climatic factors depended on the native ranges of the herbs and climatic conditions during their growing season (Körner, 2021). A thorough investigation of each specific case would be beneficial for decision making.

Some caveats need to be considered when interpreting our results. First, broader planting of the same genotype (i.e., covering all provinces in South China and even some Southeast Asian countries) would provide a thorough understanding of the optimal climatic variables resulting in the greatest consistency and efficacy of the medicinal components of camphor tree (Sharma et al., 2020). Second, the further experimental verification of the effect of temperature on metabolite contents and the functional validation of selected candidate genes (such as *TPS-b*, *TPS-g*, of *TPS-a*) of the biosynthetic pathway under controlled laboratory conditions would enhance our conclusions (Lau and Sattely, 2015; Nett et al., 2020). Third, soil characteristics (including nutritional status, humidity and rhizosphere microorganisms, etc.) are crucial for the accumulation of secondary metabolites (Ciancio et al., 2019; de Vries et al., 2020). However, owing to the absence of appropriate data for assessing these characteristics, we overlooked ecological factors such as soil conditions. Further studies addressing the abovementioned limitations would provide an in-depth framework for understanding the genetic and environmental factors related to top-geoherbalism.

Taken together, our results lay a theoretical foundation for the optimal production of this economically significant tree species. The distributional range of camphor tree largely overlaps with the land and maritime portions of the silk road of the Belt and Road Initiative (Hinsley et al., 2020); thus, products obtained from camphor tree, such as TCMs or essential oil, show a high chance of being exported to connected countries in the Middle West, Western Europe and even North Africa. As tissue culture methods for camphor trees are well established and the growth rate of the trees is relatively fast (Shi et al., 2010), the commercial cultivation of trees *ex situ* is probably a sustainable and economic method in addition to importation (Brinckmann, 2015). Thus, our research will provide guidance for policy making, genotype selection and the optimization of climatic conditions for growth by domestic and international government stakeholders, farmers, merchandisers and consumers.

## Methods

### Plant materials

The sequences utilized for *de novo* genome assembly of the genome were obtained from the fresh leaves of a single camphor tree (*C. camphora* var. *linaloolifera* Fujita; NO.95), grown at Guangxi Forestry Research Institute.

Tissue-cultured seedlings were planted in four locations in Guangxi Province in China in May 2018, including Qinzhou (21°58’52”N, 108°39’14”E), Nanning (22°49’00”N, 108°19’39”E), Baise (23°54’09”N, 106°37’06”E) and Liuzhou (24°19’35.0”N, 109°25’41”E). Fresh leaves were collected from the four locations for RNA-Seq analyses and volatile compound quantification in November 2020.

### DNA sequencing

High-molecular-weight (HMW) genomic DNA was extracted using a DNeasy Plant Mini Kit (Qiagen, USA), and 50 μg of HMW genomic DNA was prepared to generate SMRTbellTM libraries with a 20 kb insert size. Subsequently, PacBio circular consensus sequencing (CCS) data were produced on the PacBio Sequel II platform. Hi-C libraries were constructed from the tender leaves of *C. camphora* with fragments labelled with biotin and sequenced on the Illumina NovaSeq 6000 platform.

### RNA sequencing

Total RNA was extracted from each sample by using an RNAprep Pure Plant kit (TIANGEN, Beijing, China) according to the user manual. cDNA was synthesized from 20 μg of total RNA using Rever Tra Ace (TOYOBO, Osaka, Japan) with oligo (dT) primers following the user manual. High-throughput sequencing was then performed on the Illumina NovaSeq 6000 platform.

### *De novo* genome assembly and quality assessment

The *C. camphora* genome was assembled by integrating the sequencing data obtained with PacBio CCS technology and the Hi-C method using Hifiasm (Cheng et al., 2021) with the default parameters. We assembled two contig-level genomes, including a monoploid genome (haplotype A) and an allele-defined genome (haplotype B). Before Hi-C scaffolding, we filtered the redundant contigs from the contig-level genomes by using Purge_haplotigs (Roach et al., 2018). The Hi-C reads were assessed with the HiC-Pro program (Servant et al., 2015) and uniquely mapped to the contig assemblies. Juicer tools (Durand et al., 2016) and 3D-DNA pipelines (Dudchenko et al., 2017) were used to perform chromosome scaffolding. The BUSCO (Seppey et al., 2019) method was used to evaluate the completeness of the chromosome-level genomes.

### Repetitive sequences and gene annotation

We used the EDTA pipeline (Ou et al., 2019) to annotate transposable elements in the *C. camphora* genome, including LTR, TIR and nonTIR elements. LAI assessment and LTR insertion time estimation were performed by LTR_retriever (Ou and Jiang, 2018) with the synonymous substitution rate set as 3.02e-9 (Cui et al., 2006). TRF software (Benson, 1999) with modified parameters (“trf 1 1 2 80 5 200 2000”) was used to identify tandem repeats.

For gene model annotation, we trained *ab initio* gene predictors, including AUGUSTUS (Stanke and Morgenstern, 2005) and SNAP (Korf, 2004), on the repeat-masked genome using a combination of protein and transcript data. We used the annotated proteome data of *A. thaliana, L. chinense* (Chen et al., 2019), *C. kanehirae* (Chaw et al., 2019) and *P. americana* (Rendón-Anaya et al., 2019) and the Swiss-Prot database as the protein data. We employed RNA-Seq data from the four locations as the transcript data. To train AUGUSTUS, BRAKER2 (Bruna et al., 2021) was applied with the transcript data from aligned RNA-Seq bam files produced by Hisat2 (Kim et al., 2015). SNAP was trained under MAKER (Cantarel et al., 2008) with two iterations. Transcript data were supplied in the form of a *de novo* assembled transcriptome generated in Trinity (Haas et al., 2013) and a reference-based assembly generated by StringTie (Pertea et al., 2016). After training, the AUGUSTUS and SNAP results were fed into MAKER again along with all other data to produced synthesized gene models.

Functional annotations of the protein-coding sequences were obtained via BLASTP (“-evalue 1e-10”) searches against entries in both the NR and Swiss-Prot databases. The prediction of gene sequence motifs and structural domains was performed using InterProScan (Jones et al., 2014). The annotations of the GO terms and KEGG pathways of the genes were obtained from eggNOG-mapper (Huerta-Cepas et al., 2017).

### Phylogenetic analyses and estimation of divergent times

A total of 16 plant species, including five magnoliids (*C. camphora*, *C. kanehirae*, *P. americana*, *Ch. salicifolius* and *L. chinense*), four monocots (*Zea mays, Oryza sativa, Ananas comosus* and *Musa acuminata*), five eudicots (*A. thaliana*, *Populus trichocarpa*, *Medicago tuncatula*, *Solanum lycopersicum* and *V. vinifera*) and two ANA-grade species *(Nymphaea colorata* and *Amborella trichopoda*) were used to infer the phylogenetic tree. All genomes except for that of *Ch. salicifolius* (http://xhhuanglab.cn/data/Chimonanthus_salicifolius.html) were downloaded from JGI (ftp://ftp.jgi-psf.org/pub/compgen/phytozome/v12.0/) and Ensembl Plants (http://plants.ensembl.org/info/data/ftp/index.html). Paralogues and orthologues were identified among the 16 species by using the OrthoFinder pipeline (Emms and Kelly, 2019) with the default parameters, and the protein sequences of the 172 identified single-copy orthologous genes were used for the construction of the phylogenetic tree. The concatenated amino acid sequences were aligned with MAFFT (Katoh and Standley, 2013) and trimmed with trimAI (Capella-Gutiérrez et al., 2015). A maximum likelihood (ML) phylogenetic tree was constructed using IQ-TREE (Nguyen et al., 2015) with ultrafast bootstrapping (-bb 1000), using *N. colorata* and *A. trichopoda* as outgroups. The best-fitting substitution models were selected by ModelFinder (Kalyaanamoorthy et al., 2017). In addition, ASTRAL-III v5.7.3 (Zhang et al., 2018) was applied to infer the coalescence-based species tree with 172 gene trees. The species tree was then used as an input to estimate the divergence time in the MCMCTree program in the PAML package (Yang et al., 2007). The calibration time was obtained from TimeTree (Kumar et al., 2017): 107-135 Mya for the divergence of *V. vinifera* and *A. thaliana*, 42-52 Mya for *Z. mays* and *O. sativa*, 97-116 Mya for *O. sativa* and *M. acuminata* and 173-199 Mya for *A. trichopoda* and *A. thaliana*. The expansion and contraction of gene families were inferred with CAFÉ (De Bie et al., 2006) based on the chronogram of the 16 species

### Genome duplication and syntenic analyses

To identify the pattern of genome-wide duplications in *C. camphora*, we divided duplicated genes into five categories, WGD, TD, PD (duplicated genes separated by less than 10 genes on the same chromosome), TRD, and DSD (the remaining duplicates other than the four specified types) gene, using DupGen_Finder (Qiao et al., 2019) with the default parameters. The *Ka, Ks*, and *Ka/Ks* values were estimated for duplicated gene pairs based on the YN model in *KaKs*_Calculator2 (Wang et al., 2010), followed by the conversion of amino acid alignments into the corresponding codon alignments with PAL2NAL (Suyama et al., 2006). The genes in the five duplicate categories were further subjected to GO and KEGG analyses with the R package clusterProfiler (Yu et al., 2012). The enriched items were selected according to an FDR criterion of 0.05. The dating of ancient WGDs and orthologue divergence were estimated based on the formula T = *Ks*/2*r*, where *Ks* refers to the synonymous substitutions per site, and *r* (3.02e-9) is the synonymous substitution rate for magnoliids (Cui et al., 2006).

Genomic synteny blocks to be employed for intra- and interspecies comparisons among magnoliids were identified with MCscan software (Tang et al., 2008). We performed all-against-all LAST analyses (Kielbasa et al., 2011) and chained the LAST hits according to a distance cut-off of 10 genes, requiring at least five gene pairs per synteny block. The syntenic “depth” function implemented in MCscan was applied to estimate the duplication histories of the respective genomes. Genomic synteny was visualized with the Python version of MCScan (Tang et al., 2008), the R package RIdeogram (Hao et al., 2020) and Circos (Krzywinski et al., 2009).

### Gene expression profiling

The raw RNA-Seq data were filtered by using FASTP (Chen et al., 2018). The clean data were aligned to our assembled *C. camphora* genome with Hisat2 (Kim et al., 2015), and the quantification of gene expression was calculated with StringTie (Pertea et al., 2015). The Python script preDE.py built into StringTie was used to convert the quantification results into a count matrix. DEGs were detected with the DESeq2 R package (Love et al., 2014) with an FDR < 0.05.

### *TPS* gene family

*TPS* genes are classified into seven subfamilies, including *TPS-a*, *TPS-b*, *TPS-c*, *TPS-d*, *TPS-e*, *TPS-f* and *TPS-g* (Chen et al., 2011). The *TPS* genes included in the *TPS-a* subfamily are responsible for forming 15-carbon sesquiterpenoids. The *TPS-b* and *TPS-g* superfamilies encode the enzymes producing 10-carbon monoterpenoids. The *TPS-c, TPS-e* and *TPS-f* subfamilies encode diterpene synthases, which catalyse the formation of 20-carbon isoprenoids. The *TPS-d* subfamily is gymnosperm specific and encodes enzymes involved in the production of 20-carbon isoprenoids (Martin et al., 2004). The *TPS* genes were predicted based on both their conserved domains (PF01397 and PF03936) and BLAST analyses. Conserved domains were used as search queries against the predicted proteome using hmmsearch in HMMER (https://www.ebi.ac.uk/Tools/hmmer/). *TPS* protein sequences from *A. thaliana* and *C. kanehirae* were used as queries to identify the *TPS* genes of *C. camphora*, *P. americana*, *L. chinense* and *V. vinifera*. The protein sequence hits of *TPS* genes were aligned with MAFFT (Katoh and Standley, 2013) and trimmed with trimAI (Capella-Gutierrez et al., 2009). The *TPS* gene tree was constructed using RAxML (Stamatakis, 2014) with 1,000 bootstrap replicates. The *TPS-c* subfamily was designated as the outgroup. The distribution of *TPS* genes on the chromosomes was visualized in TBtools (Chen et al., 2020).

### Identification of genes involved in the MVA and MEP pathways

The MVA pathway involves six gene families, including the *acyl-coenzyme A-cholesterol acyl-transferase* (*ACAT*), *hydroxymethylglutaryl coenzyme A synthase* (*HMGS*), *hydroxymethylglutaryl coenzyme A reductase* (*HMGR*), *mevalonate kinase* (*MVK*), *phospho-mevalonate kinase* (*PMK*), and *mevalonate diphosphate decarboxylase* (*MVD*) genes. Seven gene families are involved in the MEP pathway, including the *1*-*deoxy*-*D*-*xylulose 5*-*phosphate synthase* (*DXS*), *1-deoxy-D-xylulose 5-phosphate reductoisomerase* (*DXR*), *2*-*C*-*methyl*-*D*-*erythritol*-*4*-*phosphate cytidylyltransferase* (*MCT*), *4*-(*cytidine-5-diphospho*)-*2*-*C*-*methyl*-*D*-*erythritol kinase* (*CMK*), *MDS*, (*E*)-*4*-*hydroxy*-*3*-*methyl*-*but*-*2*-*enyl*-*pyrophosphate synthase* (*HDS*) and (*E*)-*4*-*hydroxy*-*3*-*methyl*-*but*-*2*-*enyl*-*pyrophosphate reductase* (*HDR*) genes. The genes in both the MVA and MEP pathways are well documented in the model plants. To identify candidate genes related to the two pathways in the *C. camphora* genome, we collected the protein sequences of genes in the MVA and MEP pathways identified in *A. thaliana*. Using each *A. thaliana* gene as a query sequence, BLASTP (“-evalue 1e-10”) analyses was performed to identify orthologous genes in *C. camphora*.

### Determination of volatile metabolites

Six biological replicates were sampled in each location. The samples were ground into powder in liquid nitrogen. Two grams of the powder was transferred to a 20 ml headspace vial. The vials were sealed using crimp-top caps with TFE-silicone headspace septa. In solid-phase microextraction (SPME) analyses, each vial was placed at 60°C for 12 min, and a 65 μm divinylbenzene/carboxene/polydimethylsilioxane fibre (Supelco, Bellefonte, PA, USA) was then exposed to the headspace of the sample for 30 min at 60°C. Volatile metabolites were detected by MetWare (http://www.metware.cn/) based on the Agilent 7890B-7000D platform. The desorption of the volatile metabolites from the fibre coating was carried out in the injection port of the GC apparatus at 250°C for 10 min in spitless mode. The identification and quantification of volatile metabolites were carried out with a 30 m × 0.25 mm × 1.0 μm DB-5MS (5% phenyl-polymethylsiloxane) capillary column. Helium was used as the carrier gas at a linear velocity of 1.0 ml/min. The oven temperature was programmed to increase from 40°C (6 min) at 5°C/min until it reached 280°C, where it was held for 5 min. Mass spectra were recorded in electron impact (EI) ionization mode at 70 eV. The quadrupole mass detector, ion source and transfer line temperatures were set at 150, 230 and 280°C, respectively. Volatile compounds were identified by comparing the mass spectra with the data system library (MWGC) and linear retention index.

### Climatic factor collection and analyses

Data on 17 climatic factors from Qinzhou, Nanning, Baise and Liuzhou from 2018 to 2020 were downloaded from the National Meteorological Information Centre (http://data.cma.cn/en). The analyses of the climatic factors were performed in the R package FactoMineR (Lê et al., 2008).

## Funding information

This study was supported by the National Key Research and Development Program of China (Grant No. 2020YFA0907900), Genomic Studies on the Differences in the Secondary Metabolism of Camphor Tree under Different Site conditions (Grant No. JA-20-04-07 and Grant No. 202102), the National Natural Science Foundation of China (Grant No. 32070242), the Shenzhen Science and Technology Program (Grant No. KQTD2016113010482651), special funds for science technology innovation and industrial development of Shenzhen Dapeng New District (Grant No. RC201901-05 and Grant No. PT201901-19), the China Postdoctoral Science Foundation (Grant No. 2020M672904), the Basic and Applied Basic Research Fund of Guangdong (Grant No. 2020A1515110912) and the USDA Agricultural Research Service Research Participation Program of the Oak Ridge Institute for Science and Education (ORISE) (Grant No. DE-AC05-06OR23100).

## Conflict of interest

No conflict of interest was declared.

## Author contributions

L.W., C.L. K.X.L. and R.H.J. conceived and designed the study; R.H.J., X.L.C., C.S.Z., P.W. and K.X.L. prepared the materials; C.L., X.L.C., X.Z.L., D.P. and X.X.H. performed data analyses; R.H.J., X.L.C., D.E.H, C.L. and L.W. wrote the manuscript; all authors read and approved the final draft.

## Data availability

All the raw sequence reads of *C. camphora* have been deposited in NCBI under the BioProject accession number PRJNA761572. The *C. camphora* genome assembly and annotations have been deposited on the cyVerse platform (https://data.cyverse.org/dav-anon/iplant/home/licheng_caas/C.camphora_genome).

## Notes

### Competing Interest Statement

The authors have declared no competing interest.

