## Supplemental figures for "The genome of the camphor tree and the genetic and climatic relevance of the top-geoherbalism in this medicinal plant"

### Slide 1
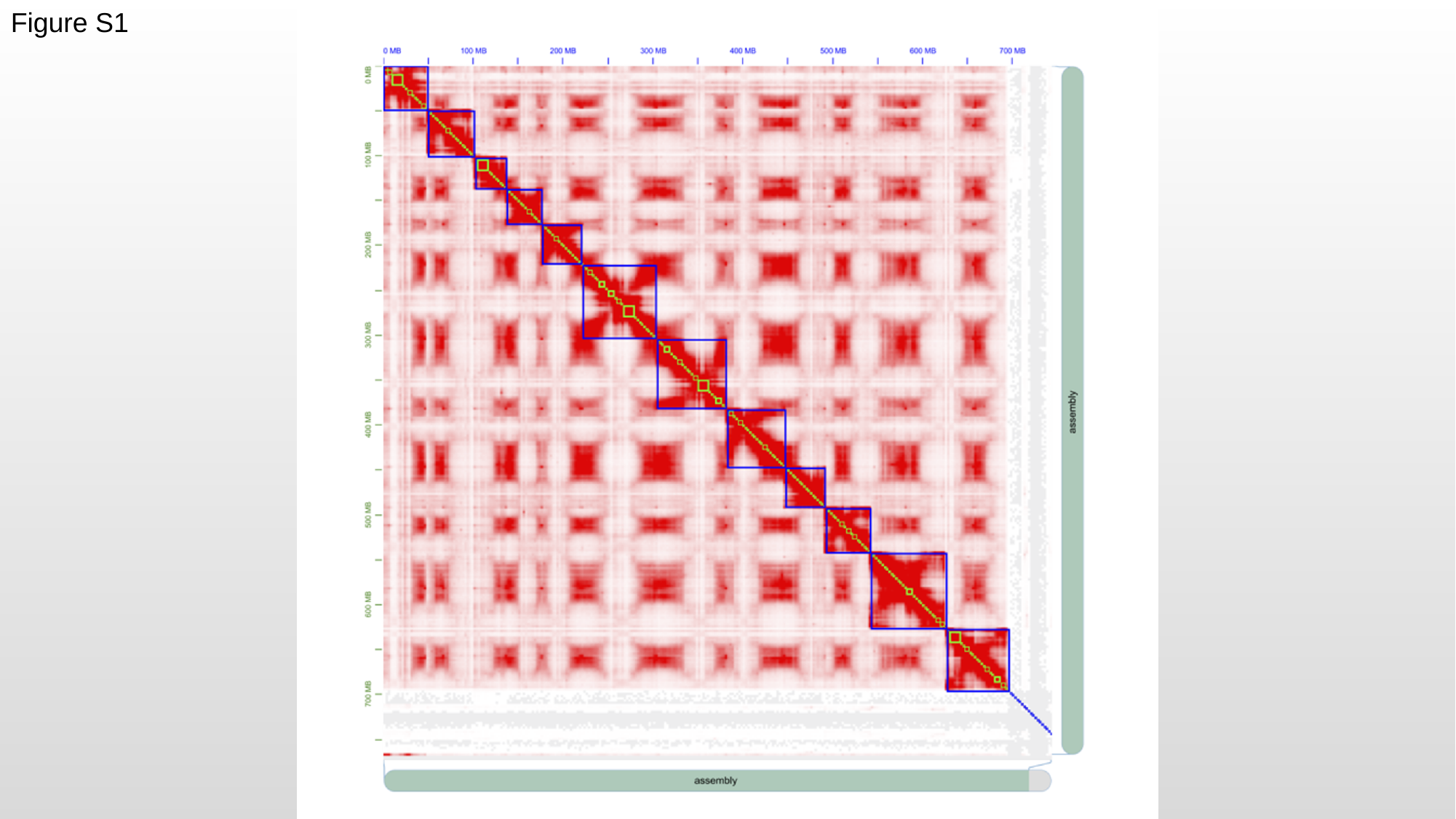

Figure S1

### Slide 2
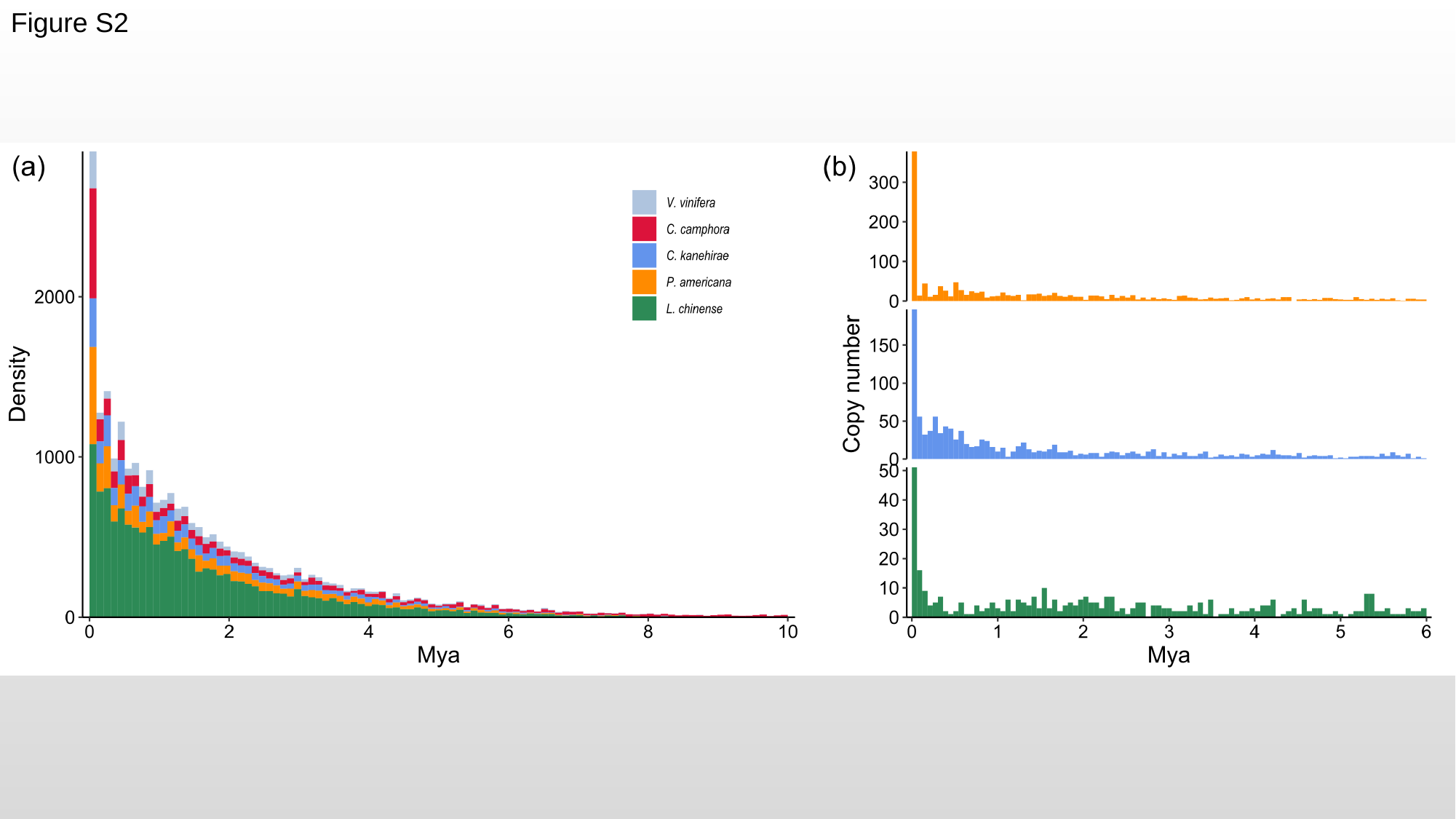

Figure S2

### Slide 3
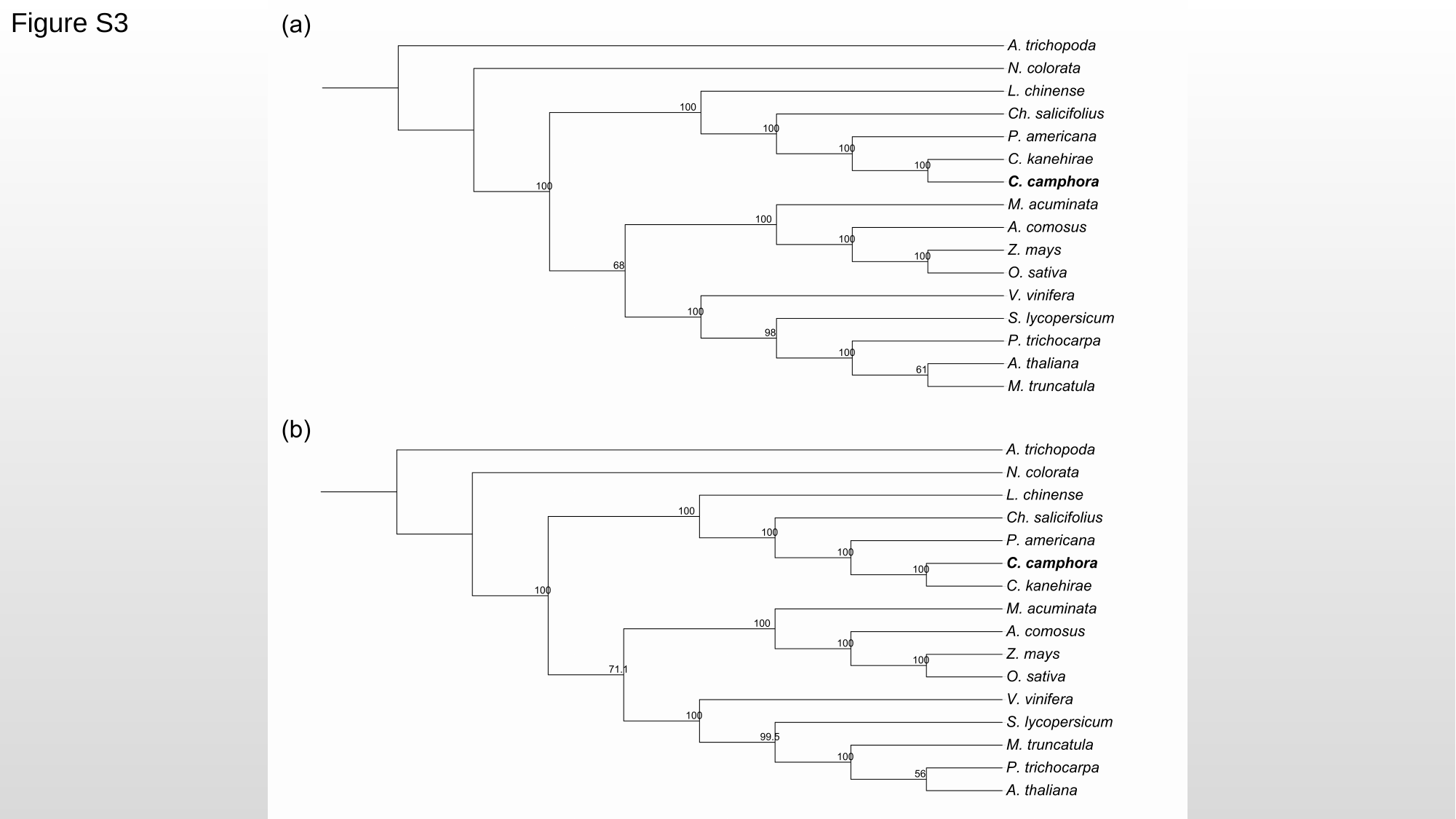

Figure S3

### Slide 4
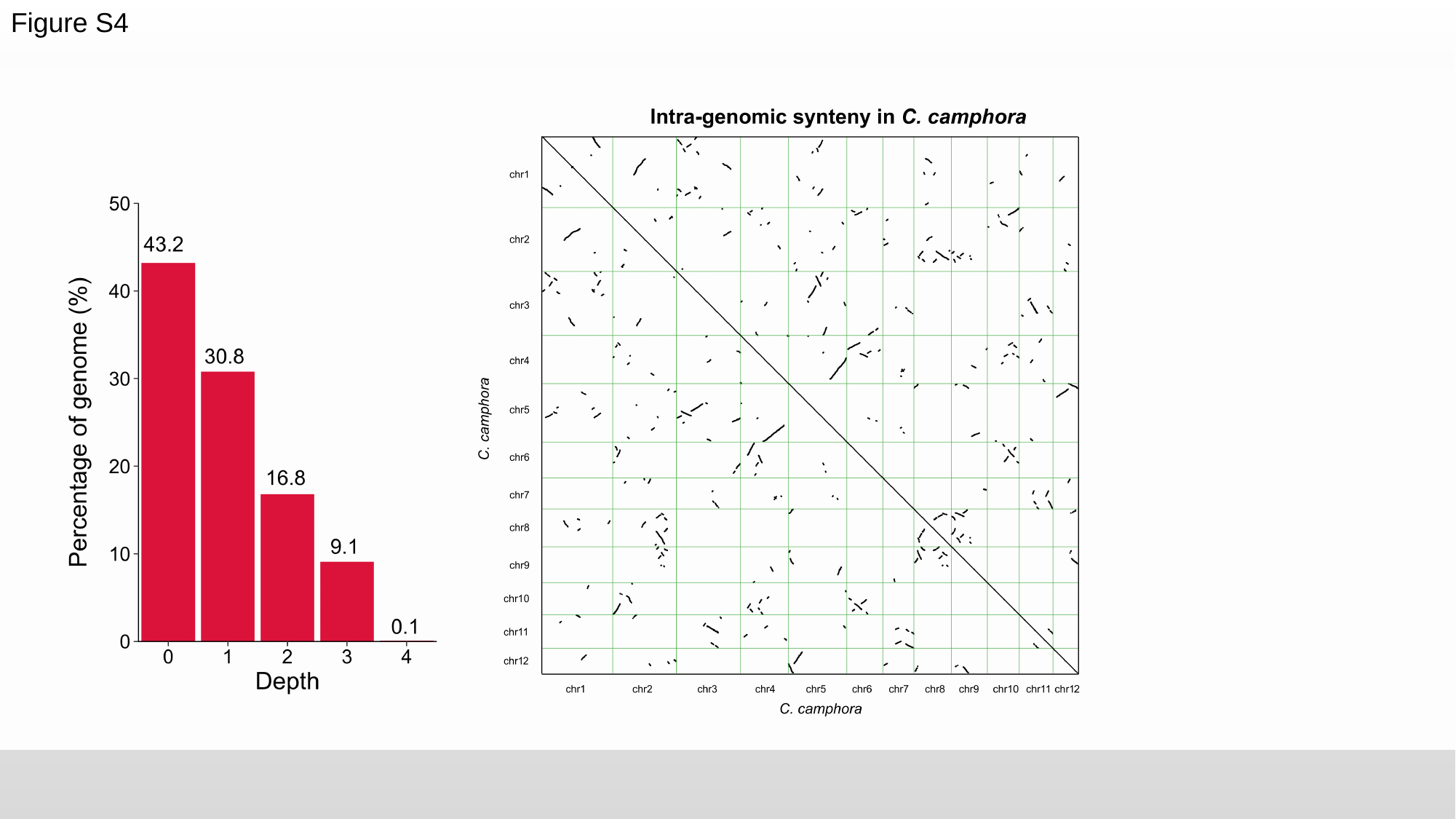

Figure S4

### Slide 5
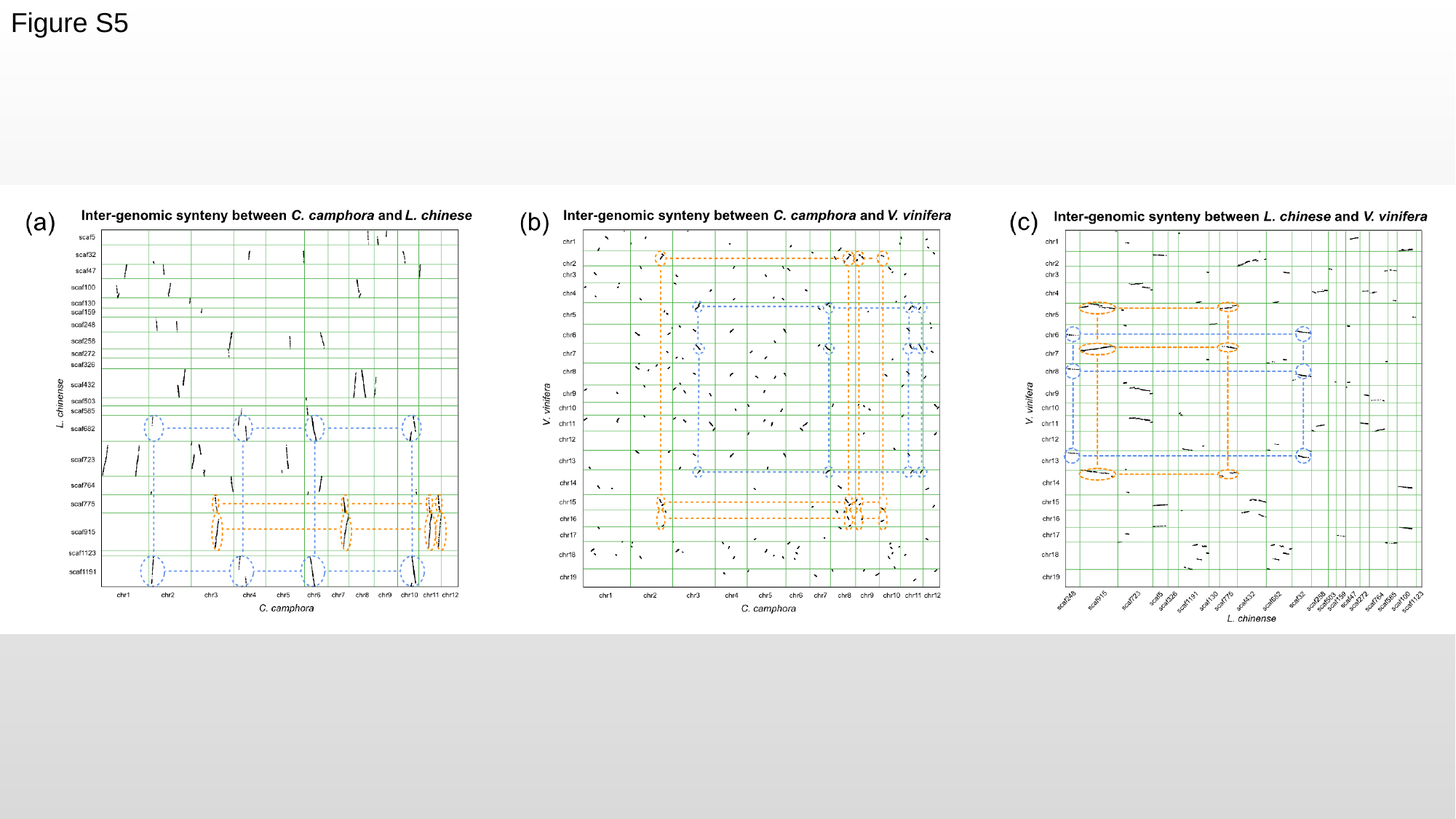

Figure S5

### Slide 6
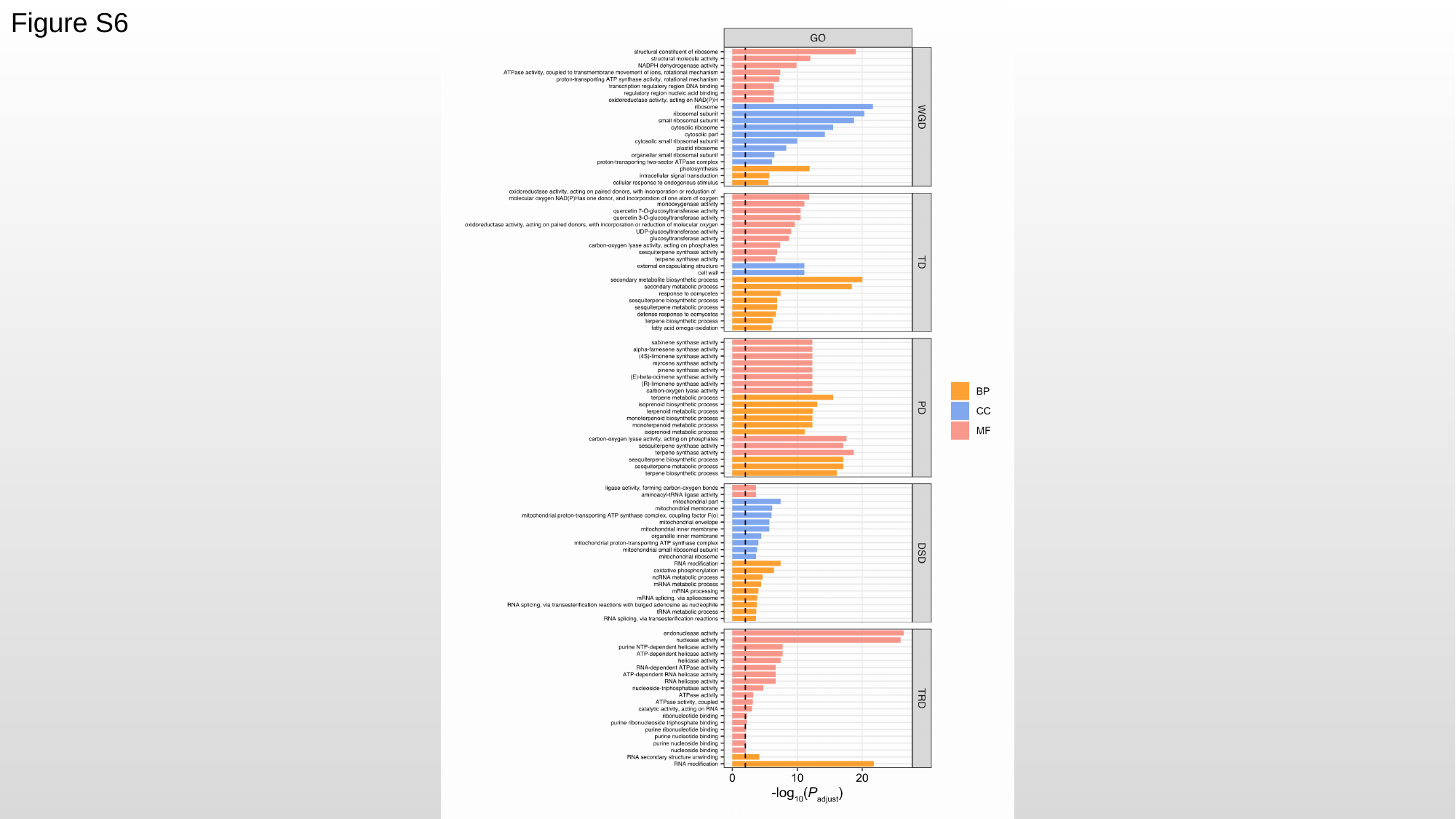

Figure S6

### Slide 7
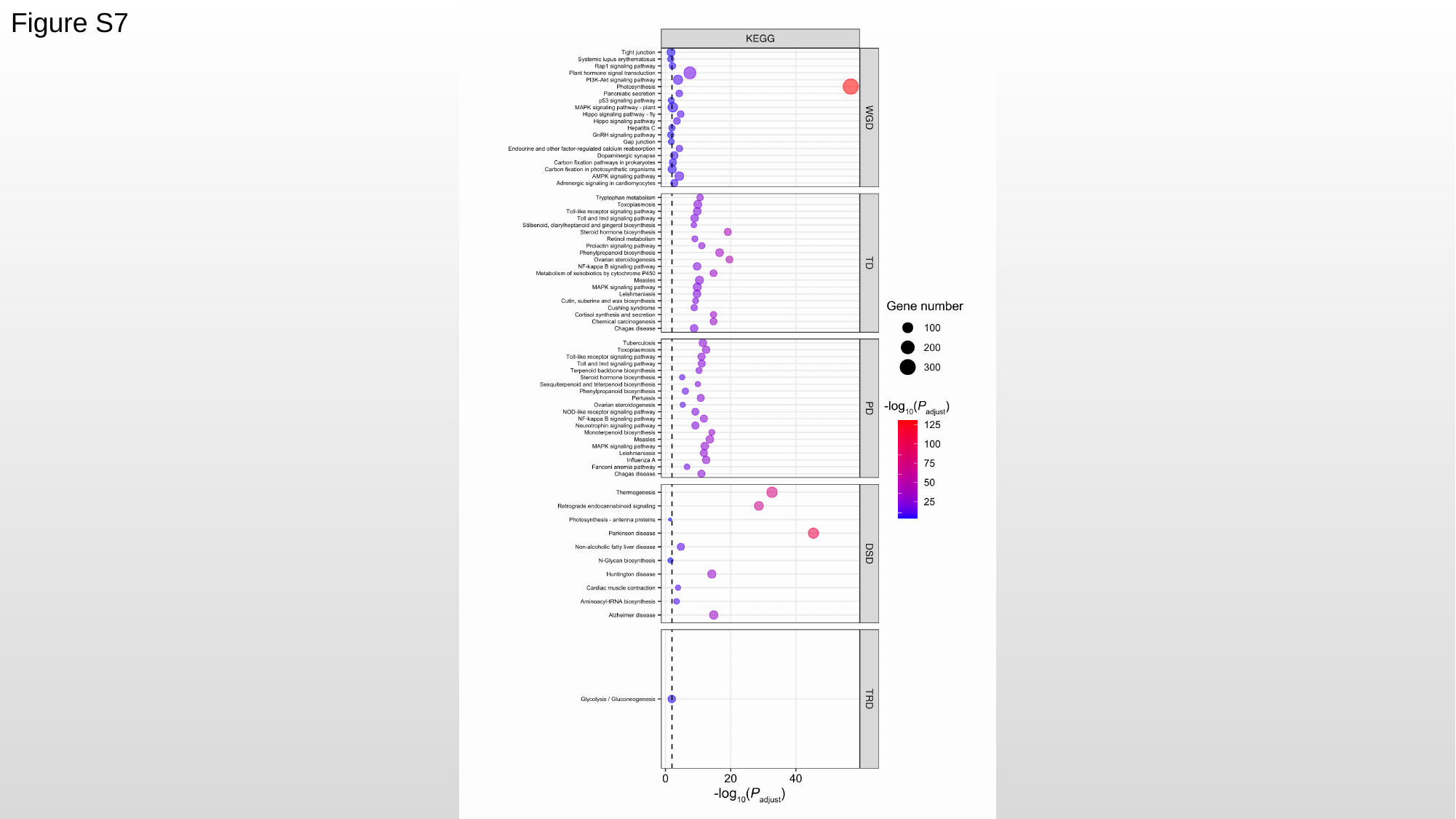

Figure S7

### Slide 8
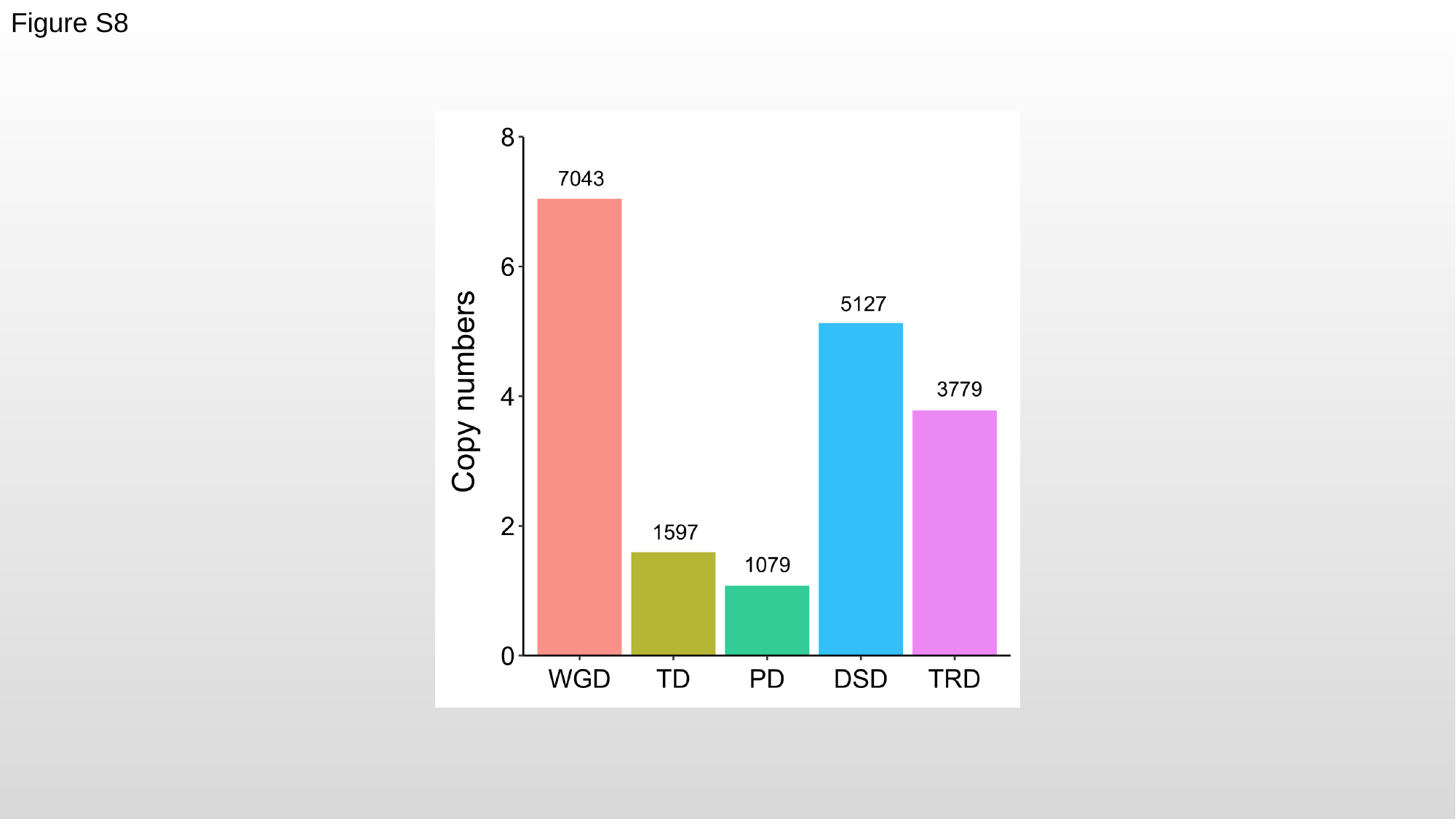

Figure S8

### Slide 9
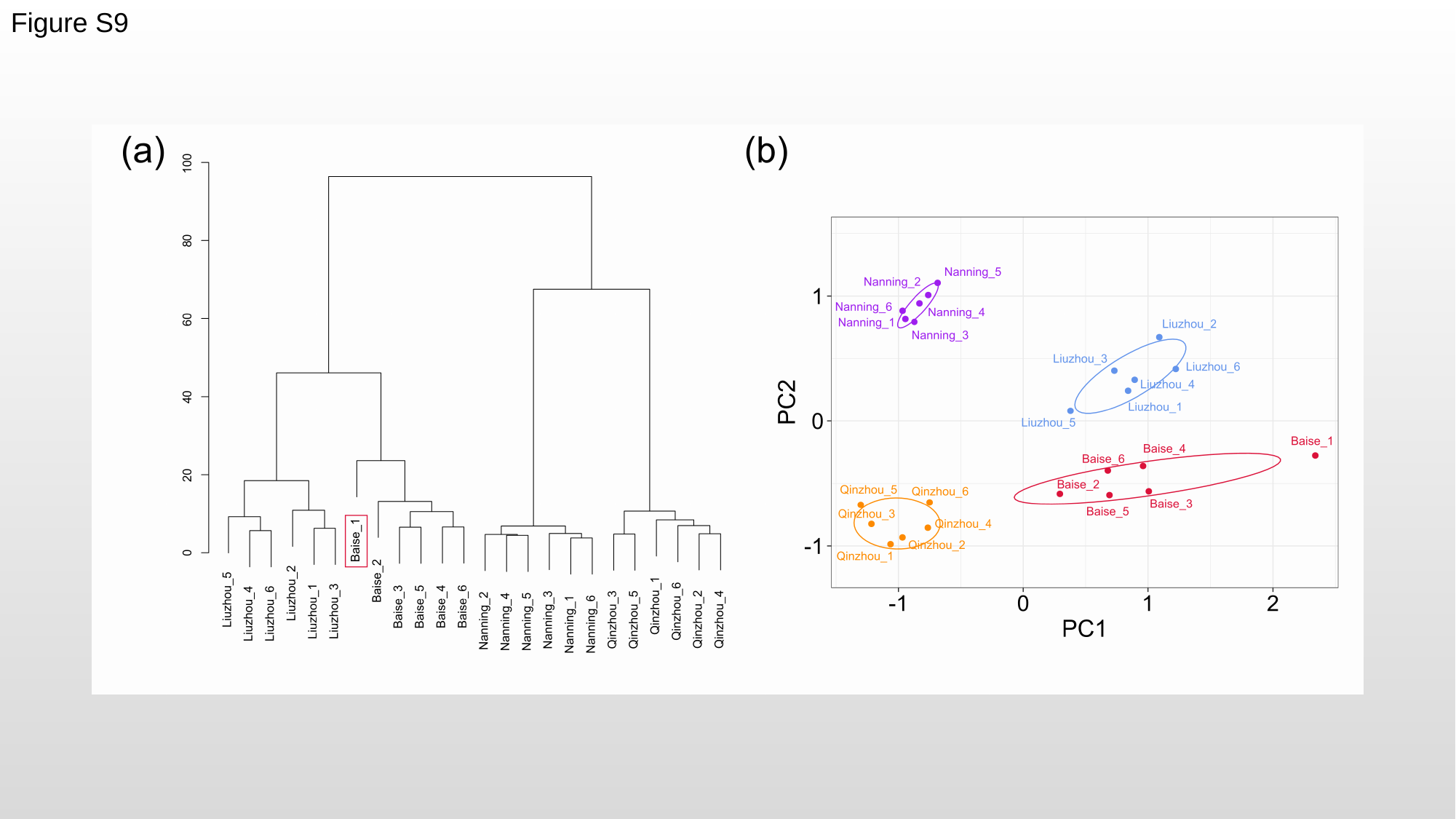

Figure S9

### Slide 10
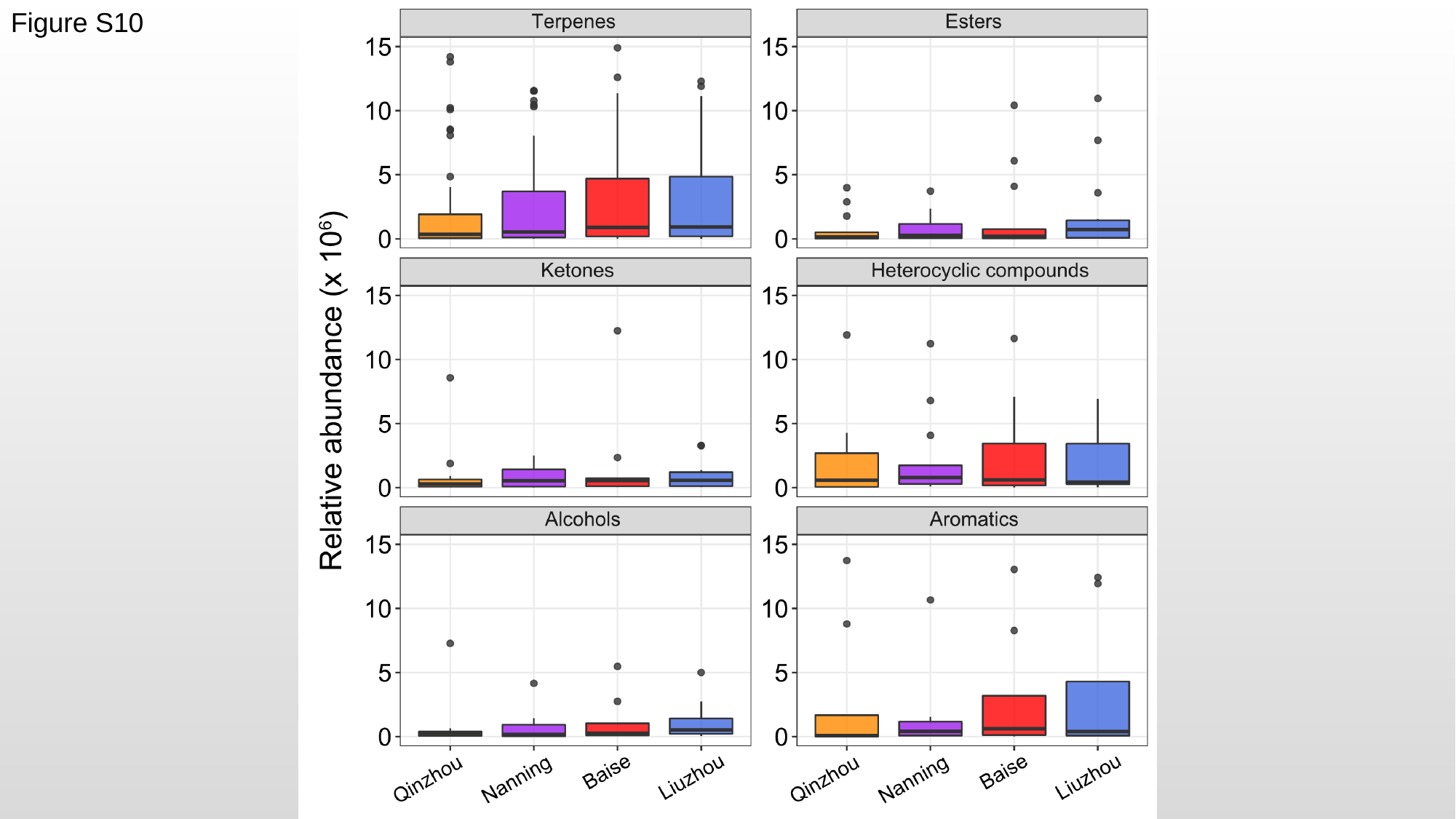

Figure S10

### Slide 11
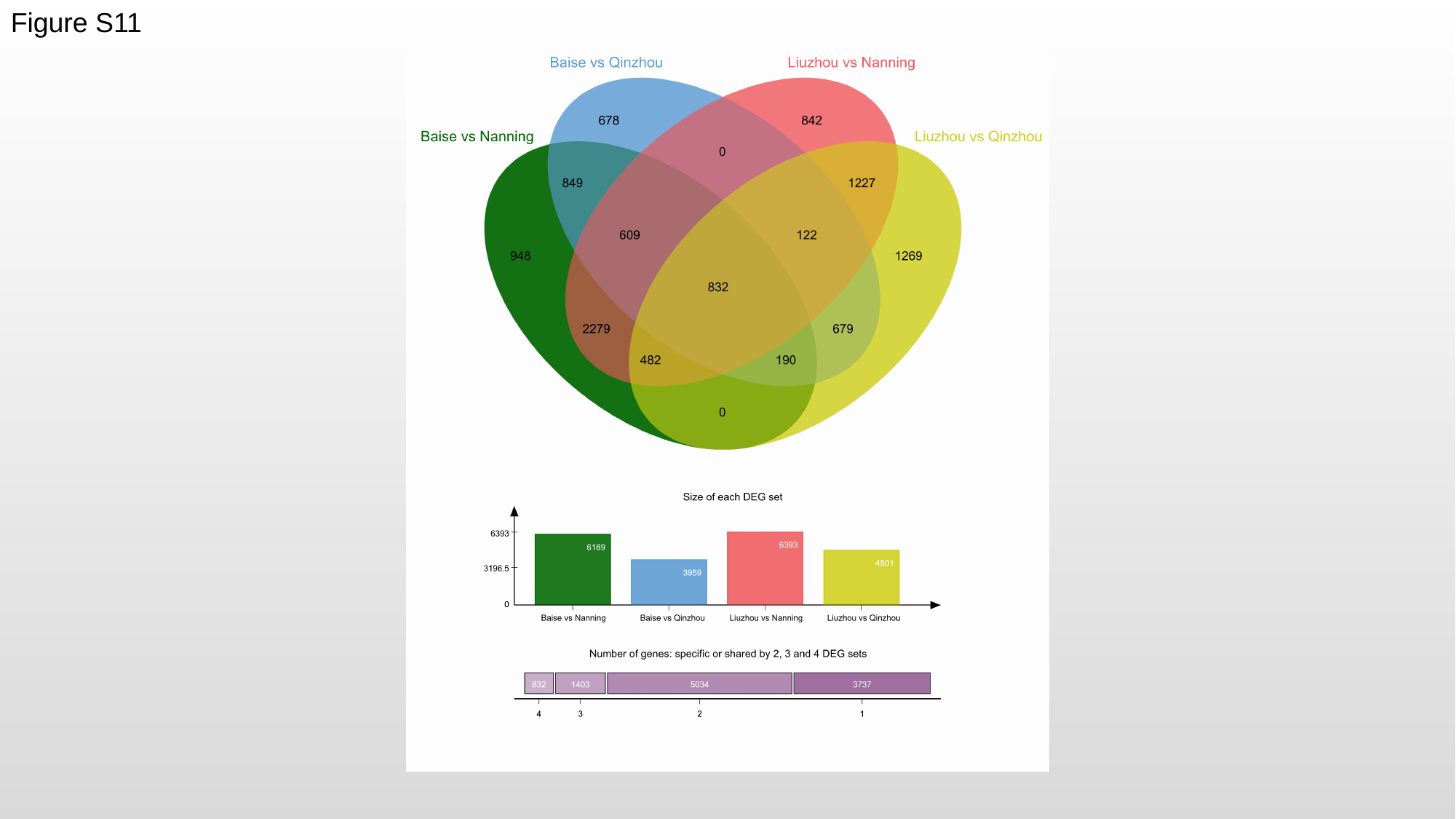

Figure S11

### Slide 12
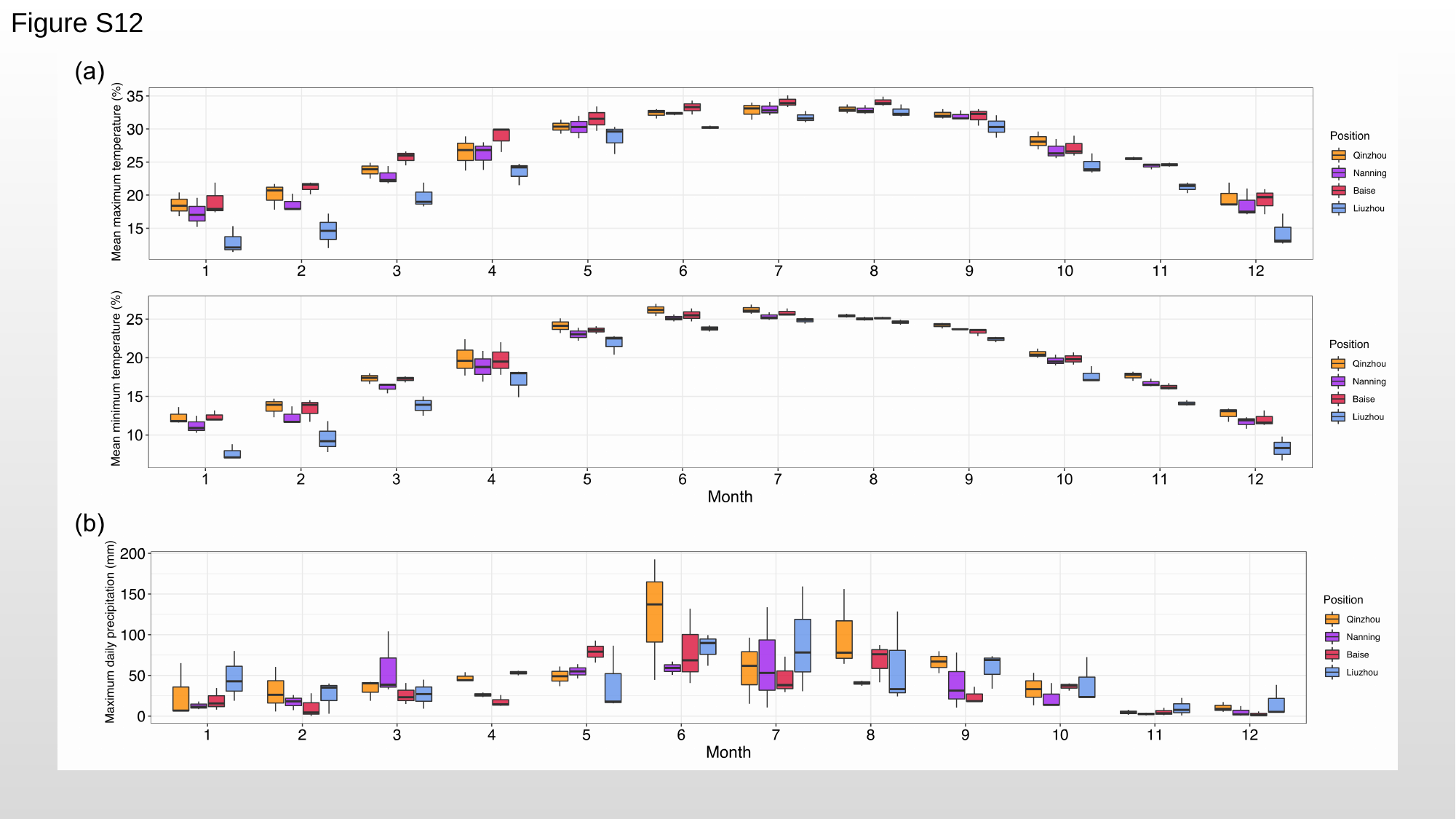

Figure S12
