## Supplemental figure legends for "The genome of the camphor tree and the genetic and climatic relevance of the top-geoherbalism in this medicinal plant"

**Figure S1** Genome-wide Hi-C heatmap of *C. camphora*. Post-clustering heatmap shows density of Hi-C interactions between contigs from 3D-DNA pipeline.

**Figure S2** Analyses of LTR insertion time. (a) Frequency distribution of LTR insertion time in magnoliids and *V. vinifera*. (b) Frequency distribution of *Copia*, *Gypsy* and unknown LTR insertion time in *C. camphora*.

**Figure S3** Reconstruction for phylogenetic trees. (a) Concatenation-based phylogenetic tree constructed by 172 strictly single-copy orthologous genes retrieved from 16 plants using IQ-TREE. (b) Coalescent-based phylogenetic tree constructed by the 172 genes. RAxML was used for construction of the 172 gene trees. The coalescent-based species tree was finally reconstructed by ASTRAL-III.

**Figure S4** Intra-genomic synteny in *C. camphora.*

**Figure S5** Inter-genomic synteny among *C. camphora*, *L. chinense* and *V. vinifera.*

**Figure S6** GO enrichment analyses of different types of duplicate genes. The abbreviations see “Methods” section.

**Figure S7** KEGG enrichment analyses of different types of duplicate genes. The abbreviations see “Methods” section.

**Figure S8** The numbers of genes among different types of duplicate genes. The abbreviations see “Methods” section.

**Figure S9** Hierarchical clustering (a) and principal component analyses (b) of metabolic abundance for quality control.

**Figure S10** The relative abundance of terpenes, esters, ketones, heterocyclic compounds, alcohols and aromatics in four different planting locations.

**Figure S11** The venn diagram and bar plot of differentially expressed genes of “Liuzhou vs Qinzhou”, “Liuzhou vs Nanning”, “Baise vs Qinzhou” and “Baise vs Nanning”.

**Figure S12** The monthly observations of mean maximum temperature, mean minimum temperature and maximum daily precipitation in the four planting locations, including Qinzhou, Nanning, Baise and Liuzhou.
